# Chemical Tools Based on the Tetrapeptide Sequence of IL-18 Reveals Shared Specificities between Inflammatory and Apoptotic Initiator Caspases

**DOI:** 10.1101/2025.02.23.639785

**Authors:** Christopher M. Bourne, Nicole R. Raniszewski, Madhura Kulkarni, Patrick M. Exconde, Sherry Liu, Winslow Yost, Tristan J. Wrong, Robert C. Patio, Ashutosh Mahale, Matilda Kardhashi, Teni Shosanya, Mirai Kambayashi, Bohdana M. Discher, Igor E. Brodsky, George M. Burslem, Cornelius Y. Taabazuing

## Abstract

Caspases are a family of cysteine proteases that act as molecular scissors to cleave substrates and regulate biological processes such as programmed cell death and inflammation. Extensive efforts have been made to identify caspase substrates and to determine factors that dictate substrate specificity. We recently discovered that that the human inflammatory caspases (caspases-1, -4, and -5) cleave the cytokines IL-1β and IL-18 in a sequence-dependent manner. Here, we report the development of a new peptide-based probe and inhibitor based on the tetrapeptide sequence of IL-18 (LESD). We found that this inhibitor was most selective and potent at inhibiting caspase-8 activity (IC_50_ = 50 nM). We also discovered that our LESD-based inhibitor is more potent than the currently used z-IETD-FMK inhibitor that is thought to be the most selective and potent inhibitor of caspase-8. Accordingly, we demonstrate that the LESD based inhibitor prevents caspase-8 activation during *Yersinia pseudotuberculosis* infection in primary bone-marrow derived macrophages. Furthermore, we characterize the selectivity and potency of currently known substrates and inhibitors for the apoptotic and inflammatory caspases using the same activity units of each caspase. Our findings reveal that VX-765, a known caspase-1 inhibitor, also inhibits caspase-8 (IC_50_ = 1 µM) and even when specificities are shared, the caspases have different efficiencies and potencies for shared substrates and inhibitors. Altogether, we report the development of new tools that will facilitate the study of caspases and their roles in biology.

## Introduction

Caspases are a family of related cysteine proteases that play key roles in regulating programmed cell death pathways such as apoptosis and pyroptosis^1-3^. Caspases are activated by diverse stimuli but are structurally related and their activation triggers proteolysis of what is estimated to be thousands of substrates to execute cell death^4-7^. Caspases that induce apoptosis are generally referred to as the apoptotic caspases, and those that induce pyroptosis are known as inflammatory caspases.

The inflammatory caspases, caspases-1, -4, -5, and -11 are activated in response to diverse pathogen-associated molecular patterns (PAMPs) or damage associated molecular patterns (DAMPs). A subset of germline-encoded pattern-recognition receptors (PRRs) detect PAMPS and DAMPS and assemble into multiprotein complexes known as inflammasomes^8, 9^. Assembly of the canonical inflammasome leads to caspase-1 dimerization, triggering its autoproteolytic activation^10-12^. Activated caspase-1 then cleaves and activates the interleukin family of cytokines, IL-1β and IL-18, and the pore-forming protein gasdermin D (GSDMD) to induce pyroptotic cell death^13-17^. In contrast to the canonical inflammasome pathway that utilizes PRRs, caspases-4/-5 in humans and caspase-11 in mice bind directly to lipopolysaccharide (LPS) from gram-negative bacteria, triggering their oligomerization and autoproteolytic activation^18^. We recently discovered that caspase-11 activation and auto processing precedes and is required to form the caspase-11 non-canonical inflammasome but whether this extends to other inflammasomes remains unknown^19^. Active caspases-4 and -5 cleave and activate IL-18 but cleave and inactivate IL-1β^20-22^. It has also been reported that active caspases-4 can cleave and activate IL-1β but this likely occurs with slower kinetics compared to the processing event that gives rise to the inactive species of IL-1β^23, 24^. Mouse caspase-11 cannot cleave IL-18 but similarly cleaves and inactivates IL-1β^20^. In this manner, the inflammatory caspases act as initiators and executioners that induce their own activation, permitting the cleavage of other downstream substrates.

Unlike the inflammatory caspases that act as both initiator and executioners, the apoptotic caspases have specialized roles and are classified into initiator (caspases-2, -8, -9, and -10) and executioner (caspases-3, -6, and -7) caspases^1, 25, 26^. Like inflammatory caspases, apoptotic initiator caspases are activated by multiprotein signaling complexes, which induce their autoproteolytic activation^27-29^. Once, activated, initiator caspases are thought to have a limited substrate repertoire, which includes the executioner caspases^4, 6, 27, 28^. Initiator caspases activate executioner caspases, which have a broader substrate profile and subsequently cleave thousands of substrates to induce apoptosis^4-6, 27, 28, 30^.

All caspases preferentially cleave peptide bonds after an aspartic acid residue in the P1 position ^31-33^. Caspases are structurally related with conserved active sites and overlapping substrate specificities^1, 3, 4, 33-38^. As a result, it has been challenging to develop selective reagents to probe the biology of specific caspases. Extensive efforts have been made to develop peptide-based reporter substrates and inhibitors, which has had limited success in producing truly selective reagents, at least using natural amino acids^39-42^. Some success has been achieved using unnatural amino acids to develop selective inhibitors of caspases-2,-8, -9, and -10, offering a promising avenue for developing selective caspase reagents^43-45^. Another strategy undertaken to develop selective caspase reagents has been to develop small molecule inhibitors, which generally relies on identifying and optimizing hits from screening a fragment-based library of covalent cysteine reactive molecules^46, 47^. This approach has led to the development of a selective caspase-6 inhibitor^48^, but specific small molecule inhibitors of other caspases remains a challenge.

Although the caspases are all structurally similar and peptide-based inhibitors and substrates display limited specificity *in vitro*, they have distinct protein substrates in cells, likely conferred by protein binding sites that are distinct from the active site known as exosites^21, 22, 49-53^. However, the molecular basis of substrate recognition by different caspases is not completely understood. We recently discovered that inflammatory caspases (caspases-1,-4,-5, and -11) recognize and cleave IL-1β and IL-18 in a sequence-dependent manner^20^. Notably, we demonstrated that substituting the amino acid recognition motif found in IL-1β (YVHD) to that found in IL-18 (LESD) facilitates IL-1β binding and activation by caspases-4 and -5, suggesting that they preferentially recognize the LESD motif. We also demonstrated that IL-18 has a stronger interaction with caspases-4 and -5 compared to caspase-1, suggesting that peptide-based inhibitors using the LESD scaffold may be specific for caspases-4 and -5^20^ and would be an invaluable tool for studying non-canonical inflammasome biology. Furthermore, apoptotic caspases have been reported to cleave and inactive IL-1β and IL-18 but whether they also recognize the tetrapeptide sequence adjacent to the cleavage site remains unclear.

Here, we designed and synthesized a new fluorogenic substrate, N-acetyl-Leu-Glu-Ser-Asp-aminomethylcoumarin (Ac-LESD-AMC), and inhibitor N-acetyl-Leu-Glu-Ser-Asp-chloromethylketone (Ac-LESD-CMK). This novel fluorogenic substrate reported on the activity of inflammatory caspases and the initiator apoptotic caspases-8 and -10, while the inhibitor potently blocked their activity. Surprisingly, the Ac-LESD-CMK inhibitor was most potent against caspase-8 and its homolog, caspase-10. Using *in vitro* kinetics, we characterized apoptotic and inflammatory caspases at the same activity units with different peptide substrates and inhibitors, revealing key insights into the selectivity profiles of caspases. We demonstrate that our new LESD inhibitor enters cells and inhibits proteolysis of inflammatory and apoptotic caspase substrates. Notably, the LESD inhibitor blocked the biologically important substrate processing mediated by caspase-8 and caspase-1 activated by *Yersinia pseudotuberculosis* infection*^54^.* Altogether, we have developed new chemical probes that will facilitate the study of both apoptotic and inflammatory caspase biology.

## Results

### Recombinant caspases cleave IL-18 and IL-1ꞵ in a tetrapeptide sequence-dependent manner

We previously reported that inflammatory caspases cleave IL-1β and IL-18 in a sequence-dependent manner,^20^ demonstrating that the LESD tetrapeptide sequence in IL-18 was recognized and processed by caspases-1, -4 and -5. Because apoptotic caspases have been reported to cleave and inactivate IL-1β and IL-18^3, 55, 56^, we first wanted to determine if the tetrapeptide motif similarly regulates the processing of IL-18 and IL-1β by apoptotic caspases. To test this, we expressed wildtype IL-18 (IL-18 WT) or IL-18 where the LESD tetrapeptide at position 33-36 was mutated to AAAD (IL-18 _33_AAAD_36_), wildtype IL-1β (IL-1β WT) or IL-1β in which the tetrapeptide sequence at position 113-116 was mutated to LESD (IL-1β _113_LESD_116_) in HEK 293T cells and incubated lysates from these cells with 0.25 activity units/μL of recombinant caspases purchased from a commercial vendor (Enzo Life Sciences) for either 1 hour or 24 hours. As previously reported, caspases-1, -4, -5, and -11 processed human IL-18 WT into the active p18 fragment while caspases-3 and -6 generated the inactivating p15 fragment (**Figure 1A**)^20, 56^. IL-18 is released by macrophages in response to infection with *Yersinia pseudotuberculosis*, which induces caspase-8 activation, and caspase-8–dependent caspase-1 activation^54, 57, 58^. Given their ability to cleave shared substrates, which of these caspases is responsible for the cleavage and release of IL-18 during *yersinia* infection has remained unclear but was presumed to be mediated by caspase-1. Consistent with this, we did not observe caspase-8 processing of IL-18 WT *in vitro* (**Figure 1A**). As anticipated, IL-18 cleavage by inflammatory caspases was tetrapeptide sequence-dependent as processing IL-18 _33_AAAD_36_ into the active p18 fragment was significantly inhibited (**Figure 1A**). Caspases-1, -3, -4, -5, and -6 processed IL-1β WT into the inactive p27 fragment and only caspase-1 was able to generate the active p17 species (**Figure 1B**). Even though IL-18 harboring the LESD sequence was only processed by inflammatory caspases, caspases-1, -4, -5, -6, -8, and -10 all processed IL-1β with the tetrapeptide sequence at position 113-116 mutated to LESD (IL-1β _113_LESD_116_) into the active p17 species after 24 h (**Figure 1B**), suggesting that some apoptotic caspases also recognize the tetrapeptide sequence but fail to cleave the native YVHD sequence of IL-1β. To confirm that processing was direct, we purified the cytokines from HEK 293T lysates and incubated them with the recombinant caspases **(Figure S1**). Consistent with the results from lysates, we observed the same processing events using purified cytokines, indicating direct processing by the caspases (**Figure S1**). These findings indicate that the tetrapeptide sequence of IL-1β regulates its processing by both inflammatory and apoptotic caspases. Curiously, although apoptotic caspases could process the LESD sequence in IL-1β, they failed to cleave this sequence in IL-18, but the reason for this currently remains unclear. This may be due to the presence of an exosite on IL-1β that is recognized by apoptotic caspases but future studies are needed to delineate the mechanistic basis of these specificities.

**Figure 1.**
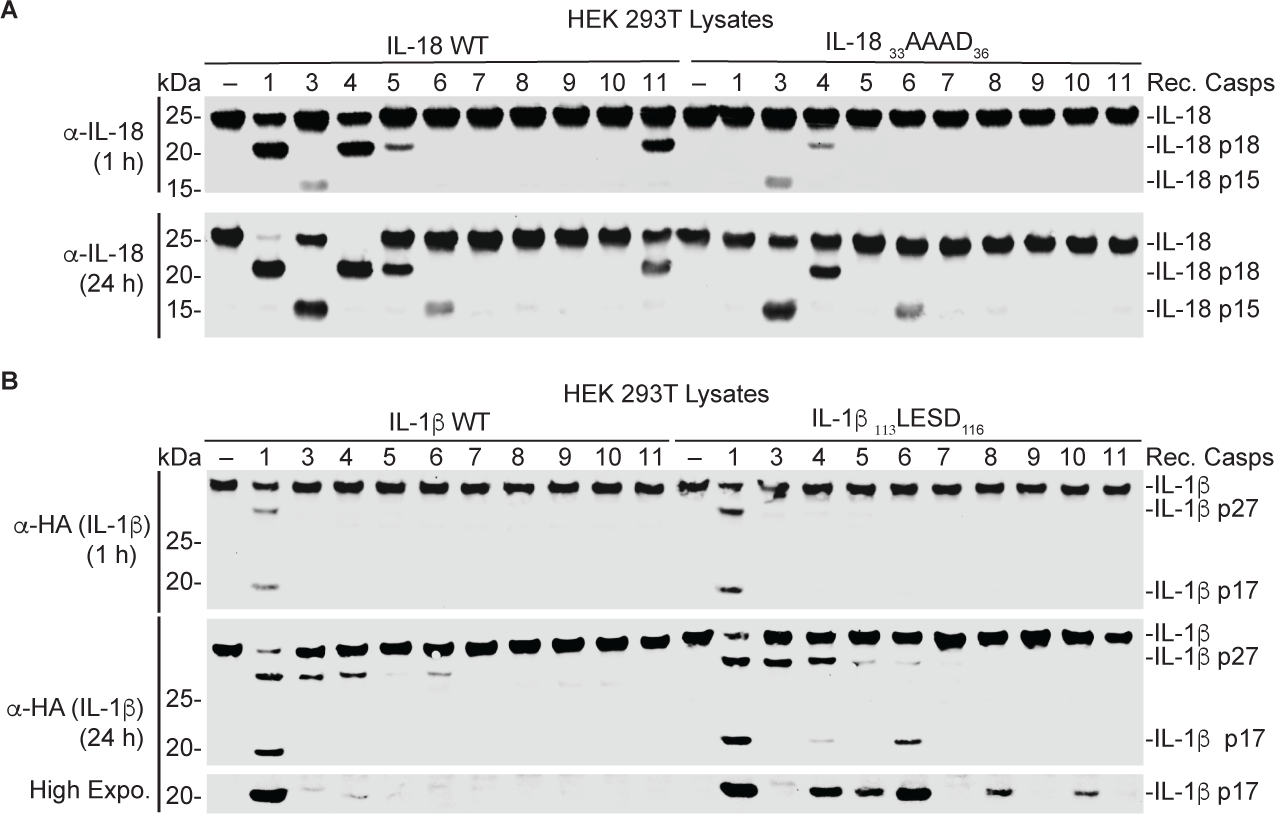
Recombinant caspases cleave IL-18 and IL-1β in a tetrapeptide sequence-dependent manner. (**A**). HA-tagged IL-18 WT and IL-18 with the tetrapeptide sequence at position 33-36 mutated to AAAD (IL-18 _33_AAAD_36_) were expressed in HEK 293T cells, and lysates were mixed with 0.25 activity units/μL of indicated caspases for 1 or 24 hours before immunoblot analysis. (**B**) IL-1ꞵ WT and IL-1ꞵ with the tetrapeptide sequence at position 113-116 mutated to LESD (IL-1β _113_LESD_116_), were expressed in HEK 293T cells and treated as in A. Data are representative of three or more independent experiments.

### An LESD-based fluorogenic substrate is selective for initiator caspases except caspase-2

The observation that replacement of the endogenous YVHD cleavage sequence in IL-1β with LESD expanded the range of caspases able to process this substrate to caspases-4, -5, -6, -8, and -10 in addition to caspase-1, suggests that a fluorogenic substrate or inhibitor based on the LESD sequence may report on the activity of these caspases. We synthesized a new fluorogenic substrate (Ac-LESD-AMC) for testing (**Figure 2A**). We incubated equivalent activity units (0.25 activity units/μL) of recombinant enzymes with the Ac-LESD-AMC substrate (**Figure 2C-K**) and used the linear rate of hydrolysis at varying substrate concentrations to generate Michaelis-Menten kinetic parameters. An example of the linear rates for caspase-1 is illustrated in **Figure 2B**. The extrinsic initiator caspases-8 and -10 (**Figure 2C-D**) and the inflammatory caspases-1, -4, -5, and -11 (**Figure 2E-H**) efficiently cleaved the Ac-LESD-AMC substrate. We did not detect processing of Ac-LESD-AMC by the executioner caspases-3, -6, and -7 (**Figure 2I-K**), suggesting that Ac-LESD-AMC is specific to initiator and inflammatory caspases. Using the kinetic parameters, we determined the relative catalytic efficiency (*k*_cat_/*K*_M_) for all caspases for Ac-LESD-AMC (**Figure 2L**). Despite no IL-18 processing *in vitro* by caspase-8, Ac-LESD-AMC was most efficiently hydrolyzed by caspase-8, which was ∼10 fold faster than the human inflammatory caspases 1, -4, -5. Caspase-10, a homolog of caspase-8, also efficiently cleaved the Ac-LESD-AMC probe. Of the caspases that cleaved the Ac-LESD-AMC probe, mouse caspase-11 was the most inefficient at cleaving Ac-LESD-AMC (∼100 fold slower than caspase-8), in agreement with the notion that caspase-11 does not cleave mouse IL-18 which harbors an LESD motif at the cleavage site. Although caspase-11 does not process mouse IL-18, it is able to process human IL-18 but less efficiently than caspase-4, and recent structural studies imply that this may be regulated by the active site sterics^20-22^. We also tested the ability of caspases to process other well-established substrates^3, 6, 32, 36, 44, 59^, including Ac-DEVD-AMC, Ac-YVAD-AMC, Ac-LEHD-AMC, and Ac-WEHD-AMC for the extrinsic initiator caspases-8 and -10 (**Figure S2-S3**), the inflammatory caspases-1, -4, -5, and mouse -11 (**Figure S4-S7**), and the executioner caspases-3, -6, and -7 (**Figure S8-10**). The kinetic values are summarized in **Table 1**.

**Figure 2.**
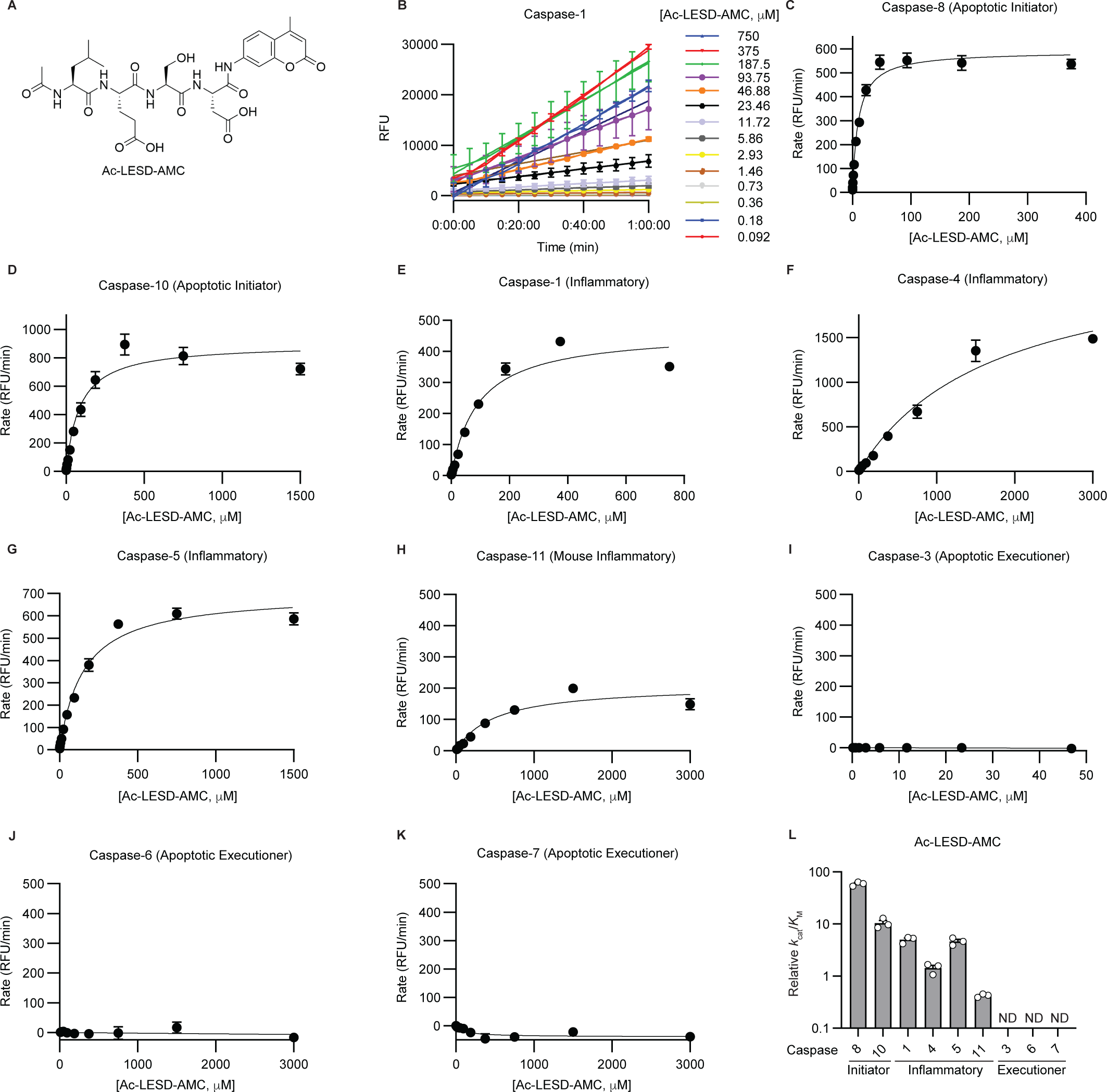
An LESD-based probe is selective for initiator caspases. (**A**) Chemical structure of the Ac-LESD-AMC fluorogenic tetrapeptide probe. (**B**) Representative linear rates of hydrolysis by 0.25 activity units/μL of caspase-1 with varying concentrations of Ac-LESD-AMC substrate. Relative fluorescence units (RFU). (**C-K**) Michaelis-Menten kinetic profiles of Ac-LESD-AMC substrate cleavage by 0.25 activity units/μL of recombinant apoptotic initiator caspases-8 and -10 (**C-D**), inflammatory caspases-1, -4, -5, and -11 (**E-H**), and apoptotic executioner caspases-3, -6, and -7 (**I-K**). Data are means ± SEM of three independent experiments with two technical replicates per experiment. (**L**) Relative catalytic efficiency, represented as *k*_cat_/*K*_M_ of caspases for Ac-LESD-AMC calculated from *in vitro* Michaelis-Menten curves. The graph depicts values calculated from three independent experiments for each caspase with the Ac-LESD-AMC probe. Data was fitted using the Michaelis-Menten equation in GraphPad Prism.

**Table 1.** Summary data for probe kinetics of each caspase with the Ac-LESD-AMC, Ac-YVAD-AMC, Ac-DEVD-AMC, Ac-LEHD-AMC, and Ac-WEHD-AMC probes. Graphs depicting the raw data are in Figures 2-3 and Figure S2-S11. *The caspase-9 preparation was different from the other caspases tested. Not assessed (N/A), not detected (ND).

|  | LESD-AMC | YVAD-AMC | DEVD-AMC | LEHD-AMC | WEHD-AMC |
| --- | --- | --- | --- | --- | --- |
| Caspase | Relative $k_{cat}/K_M$ | | | | |
| 1 | $5.01 \pm 0.41$ | $23.9 \pm 2.79$ | $1.41 \pm 0.43$ | $45.36 \pm 8.42$ | $334 \pm 48.0$ |
| 3 | ND | ND | $482 \pm 76.5$ | $4.61 \pm 0.18$ | $1.33 \pm 0.04$ |
| 4 | $1.44 \pm 0.18$ | $0.58 \pm 0.08$ | $1.27 \pm 0.18$ | $10.8 \pm 1.02$ | $10.6 \pm 0.33$ |
| 5 | $4.65 \pm 0.43$ | $7.88 \pm 0.73$ | ND | $17.6 \pm 2.00$ | $21.6 \pm 2.38$ |
| 6 | ND | ND | $13.3 \pm 2.24$ | $8.93 \pm 0.70$ | $2.28 \pm 0.31$ |
| 7 | ND | ND | $561 \pm 81.2$ | $1.07 \pm 0.17$ | $0.31 \pm 0.03$ |
| 8 | $59.2 \pm 3.26$ | $1.84 \pm 0.84$ | $310 \pm 60.4$ | $150 \pm 25.9$ | $14.3 \pm 2.92$ |
| 9* | $0.15 \pm 0.02$ | N/A | ND | $0.86 \pm 0.10$ | $0.35 \pm 0.08$ |
| 10 | $10.4 \pm 1.30$ | ND | $7.20 \pm 1.02$ | $12.3 \pm 3.06$ | $0.91 \pm 0.23$ |
| 11 | $0.42 \pm 0.02$ | ND | ND | $5.19 \pm 0.77$ | $6.97 \pm 0.68$ |
| Caspase | $V_{max}$ (RFU/min) | | | | |
| 1 | $465.55 \pm 10.25$ | $258.15 \pm 6.21$ | $44.68 \pm 15.51$ | $2702.91 \pm 85.48$ | $3496.13 \pm 121.4$ |
| 3 | ND | ND | $940.88 \pm 31.57$ | $288.98 \pm 33.2$ | $93.42 \pm 8.73$ |
| 4 | $2546.06 \pm 156.63$ | $1668.58 \pm 190.22$ | $862.73 \pm 31.55$ | $2821.37 \pm 159.35$ | $6565.47 \pm 587.59$ |
| 5 | $703.19 \pm 24.07$ | $555.56 \pm 19.58$ | ND | $1207.5 \pm 40.91$ | $1372.07 \pm 49.43$ |
| 6 | ND | ND | $455.67 \pm 64.18$ | $791.57 \pm 16.74$ | $208.71 \pm 17.47$ |
| 7 | ND | ND | $1266.91 \pm 38.79$ | $139.05 \pm 12.48$ | $32.33 \pm 4.07$ |
| 8 | $590.08 \pm 30.37$ | $19.15 \pm 4.44$ | $218.27 \pm 2.74$ | $1101.52 \pm 37.65$ | $279.84 \pm 9.36$ |
| 9* | $208.50 \pm 45.00$ | N/A | ND | $139.43 \pm 8.11$ | $38.80 \pm 1.35$ |
| 10 | $898.63 \pm 57.69$ | ND | $476.26 \pm 38.71$ | $1043.38 \pm 77.53$ | $130.96 \pm 19.73$ |
| 11 | $211.01 \pm 15.01$ | ND | ND | $1192.55 \pm 30.79$ | $1428.54 \pm 73.6$ |
| Caspase | $K_M$ (μM) | | | | |
| 1 | $93.9 \pm 6.26$ | $11.2 \pm 1.64$ | $47.5 \pm 30.8$ | $63.7 \pm 11.2$ | $10.9 \pm 1.39$ |
| 3 | ND | ND | $2.07 \pm 0.36$ | $62.4 \pm 5.40$ | $69.8 \pm 4.44$ |
| 4 | $1867 \pm 388$ | $3095 \pm 715$ | $702 \pm 76.3$ | $264 \pm 21.6$ | $626 \pm 70.6$ |
| 5 | $154 \pm 13.5$ | $72.4 \pm 9.44$ | ND | $70.1 \pm 6.28$ | $64.8 \pm 5.47$ |
| 6 | ND | ND | $35.3 \pm 5.12$ | $90.1 \pm 8.93$ | $94.7 \pm 12.7$ |
| 7 | ND | ND | $2.36 \pm 0.36$ | $138 \pm 30.0$ | $102 \pm 4.21$ |
| **8** | 9.98 ± 0.37 | ND | 0.75 ± 0.12 | 7.80 ± 1.32 | 22.0 ± 6.04 |
| **9\*** | 1513.83 ± 441.20 | N/A | ND | 167.00 ± 23.86 | 127.32 ± 36.01 |
| **10** | 87.9 ± 5.38 | ND | 70.3 ± 14.6 | 96.4 ± 23.3 | 154 ± 20.2 |
| **11** | 503 ± 58.4 | ND | ND | 242 ± 41.4 | 211 ± 33.8 |

In contrast to the other caspases, commercially purchased recombinant caspase-9 from the same supplier did not exhibit any detectable activity when incubated with fluorogenic substrates (data not shown). To overcome this limitation, we purchased full-length caspase-9 that autoproteolyzed into the p35 and p10 fragments from another supplier and incubated 1 activity unit/μL enzyme with Ac-LESD-AMC, Ac-DEVD-AMC, Ac-LEHD-AMC, and Ac-WEHD-AMC (**Figure S11A-D**). We observed that Ac-LEHD-AMC was most efficiently cleaved (relative *k*_cat_/*K*_M_= 0.86) followed by Ac-WEHD-AMC (relative *k*_cat_/*K*_M_= 0.35), then Ac-LESD-AMC (relative *k*_cat_/*K*_M_=0.15) (**Figure S11E, Table 1**). Recombinant caspase-9 did not cleave the Ac-DEVD-AMC substrate. Caspase-2 is also an initiator caspase but it has specificity for 5 amino-acid peptide substrates and is unlikely to cleave any of the tetrapeptide substrates tested^60^.

Direct comparison of the catalytic efficiencies for a given probe between caspases is challenging as each caspase was tested at an unknown concentration, normalized for activity. Nonetheless, the Ac-LESD-AMC probe was among the most efficiently cleaved substrates for caspases-8 and -10 within 3 fold of Ac-LEHD-AMC, their preferred substrate (**Figure 3A-B**, **Table 1**). Caspase-5 also efficiently cleaved Ac-LESD-AMC, within 5 fold of its preferred substrate Ac-WEHD-AMC (**Figure 3E**, **Table 1**), while the other inflammatory caspases-1, -4, and mouse caspase-11 preferred Ac-WEHD-AMC ∼10 fold or more than Ac-LESD-AMC (**Figure 3C-F**). All of the executioner caspases preferred the Ac-DEVD-AMC substrate and failed to cleave Ac-LESD-AMC (**Figure 3G-I**). Altogether our data suggests that the initiator caspases-8 and -10, and the inflammatory caspase-5, have the highest catalytic efficiency for Ac-LESD-AMC. Importantly, caspases have distinct substrate specificities and when substrate specificities overlap, the rates of hydrolysis often differ between caspases. We tested these caspases at the same activity units, allowing for direct comparison of the selectivity profile and rates of hydrolysis for given substrates. This approach has previously been utilized to offer critical insight into the relative importance of substrates for caspases^6^. Our data also suggest that other factors such as the recently reported exosites^21, 22, 49-53^, enzyme-substrate binding sites that are distinct from the active site, may play a major role in dictating protein substrate specificities as the peptide-based substrate profiles do not always recapitulate the protein substrate specificities.

**Figure 3.**
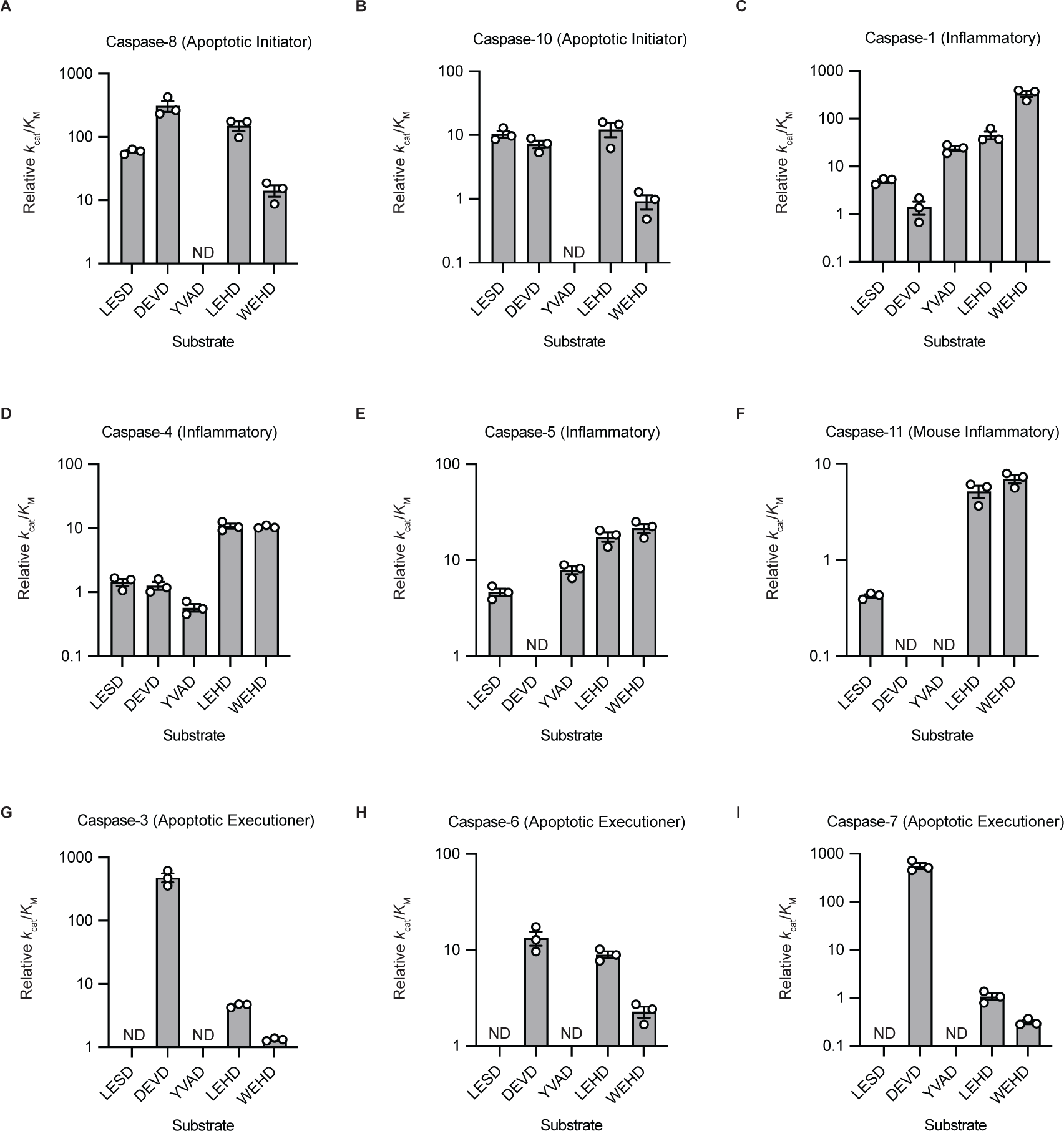
Evaluation of caspase specificities for peptide-based substrates. (**A – I**) Catalytic efficiency, represented as *k*_cat_/*K*_M_ of caspases with different tetrapeptide substrates probes. *In vitro* kinetics were fitted using the Michaelis-Menten function in GraphPad Prism to generate the kinetic parameters used to calculate *k*_cat_/*K*_M_ values for recombinant apoptotic initiator caspases-8 and -10 (**A-B**), inflammatory caspases-1, -4, -5, and -11 (**C-F**), and apoptotic executioner caspases-3, -6, and -7 (**G-I**). Data are means ± SEM of three independent experiments. ND (Not Detected).

### Selectivity profile of peptide-based caspase inhibitors

Given that the Ac-LESD-AMC substrate was most efficiently processed by caspase-8 and -10 followed by the inflammatory caspase-5, we reasoned that we could use this peptide to develop a potent inhibitor for caspase-8. We therefore synthesized the new substrate peptide with a c-terminal chloromethylketone warhead (Ac-LESD-CMK), a classical method for the inhibition of cysteine proteases, and tested the ability of this new inhibitor against commercially available caspase-8 inhibitors, z-LEHD-FMK and z-IETD-FMK, and the caspase-1/-4 inhibitor VX-765, *in vitro* (**Figure 4**)^61, 62^. The chemical structures of the inhibitors tested are depicted in **Figure 4A**. Each inhibitor was tested under saturating conditions of the most efficiently processed substrate for each specific caspase. For example, caspase-1 inhibition was assessed using Ac-WEHD-AMC as a substrate and caspase-3 inhibition was assessed using Ac-DEVD-AMC as a substrate. The IC_50_ values for each caspase and inhibitor are summarized in **Table 2**. Caspase-8 was most effectively inhibited by Ac-LESD-CMK (IC_50_ = 50 nM), followed by z-LEHD-FMK (IC_50_ = 0.70 nM), then z-IETD-FMK (IC_50_ = 350 nM) (**Figure 4B**, **Table 2**). VX-765 also inhibited caspase-8 (IC_50_ = 1 μM), despite being marketed as a caspase-1 inhibitor. Caspase-10 was also most effectively inhibited by Ac-LESD-CMK (IC_50_ = 520 nM), followed by z-LEHD-FMK (IC_50_ = 3.59 μM), then z-IETD-FMK (IC_50_ = 5.76 μM), but weakly inhibited by VX-765 (IC_50_ = 42 μM) (**Figure 4C**, **Table 2**). VX-765 was the most efficient inhibitor for caspase-1 (IC_50_ = 530 nM) (**Figure 4D**, **Table 2**). Ac-LESD-CMK was also the most efficient inhibitor of caspase-5 tested (IC_50_ = 2 μM) (**Figure 4E**, **Table 2**). Broadly, Ac-LESD-CMK and VX-765 weakly inhibited the executioner caspases, while z-LEHD-FMK and z-IETD-FMK were the two most efficient inhibitors tested for the executioner caspases (**Figure 4F-H**, **Table 2**). Taken together, these data demonstrate that Ac-LESD-CMK is a potent and selective inhibitor of caspases-8, -10, and -5. Notably, the commercially available caspase-8 inhibitor z-IETD-FMK is less potent for caspase-8 and lacks selectivity as it also blocks the activity of the executioner caspases. VX-765 inhibits caspase-1, as expected, but also blocks caspase-8 activity. z-LEHD-FMK was among the most efficient inhibitors for all caspases tested and is not a selective inhibitor.

**Figure 4.**
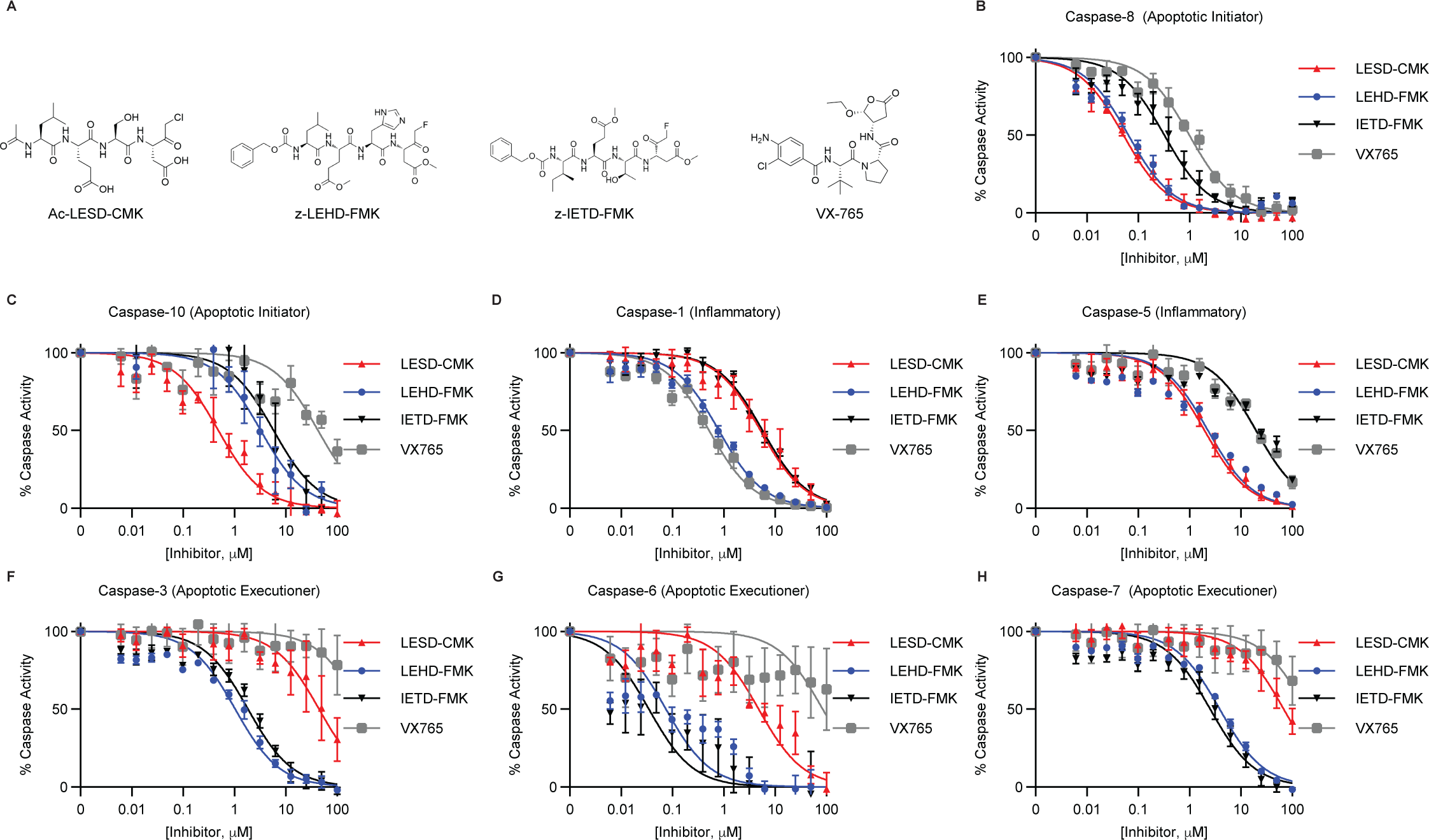
Selectivity profile of peptide-based caspase inhibitors. Each caspase was incubated with a saturating concentration of its preferred peptide substrate, in the presence of either the Ac-LESD-CMK, z-LEHD-FMK, z-IETD-FMK, or VX-765 inhibitors at indicated concentrations. Substrate cleavage rates were determined at each inhibitor concentration and normalized to the no inhibitor condition for each run. (**A**) Chemical structures of Ac-LESD-CMK, z-LEHD-FMK, z-IETD-FMK, and VX-765 inhibitors. (**B**) Caspase-8 inhibition was assessed using 200 μM Ac-LEHD-AMC substrate. (**C**) Caspase-10 inhibition was assessed using 200 μM Ac-LEHD-AMC substrate. (**D**) Caspase-1 inhibition was assessed using 200 μM Ac-WEHD-AMC substrate for activity. (**E**) Caspase-5 inhibition was assessed using 200 μM Ac-WEHD-AMC substrate. (**F**) Caspase-3 inhibition was assessed using 100 μM Ac-DEVD-AMC substrate. (**G**) Caspase-6 inhibition was assessed using 100 μM Ac-DEVD-AMC substrate. (**H**) Caspase-7 inhibition was assessed using 100 μM Ac-DEVD-AMC substrate. Data were fitted using the [Inhibitor] vs. normalized response function in GraphPad Prism. Data are means ± SEM of three independent experiments.

**Table 2.** Summary of caspase inhibitor IC_50_ values for data depicted in figures 4, S12 and S13. *The caspase-9 preparation was different from the other caspases tested. Not assessed (N/A).

|  | **LESD-CMK** | **LEHD-FMK** | **IETD-FMK** | **FLTD-CMK** | **VX-765** | **zVAD-FMK** |
| --- | --- | --- | --- | --- | --- | --- |
| **Caspase** | **IC<sub>50</sub> (μM)** |  |  |  |  |  |
| **1** | 5.67 ± 1.87 | 0.82 ± 0.13 | 5.72 ± 0.47 | 3.38 ± 0.77 | 0.53 ± 0.09 | 0.024 ± 0.006 |
| **3** | 54.0 ± 18.0 | 1.07 ± 0.18 | 1.88 ± 0.26 | 56.0 ± 27.7 | >100 | 0.030 ± 0.004 |
| **4** | 59.0 ± 5.77 | N/A | N/A | >100 | 30.1 ± 2.67 | 0.32 ± 0.011 |
| **5** | 2.03 ± 0.45 | 2.31 ± 0.31 | 19.91 ± 1.19 | 15.6 ± 1.09 | 19.5 ± 1.37 | 0.13 ± 0.011 |
| **6** | 5.08 ± 1.14 | 0.11 ± 0.06 | 0.06 ± 0.04 | 43.4 ± 15.9 | >100 | 0.08 ± 0.01 |
| **7** | 72.0 ± 12.9 | 3.90 ± 0.29 | 2.63 ± 0.54 | 53.5 ± 31.0 | >100 | 0.006 ± 0.001 |
| **8** | 0.05 ± 0.01 | 0.07 ± 0.01 | 0.35 ± 0.10 | 5.22 ± 2.35 | 1.02 ± 0.19 | 0.002 ± 0.0003 |
| **9\*** | 12.13 ± 7.64 | 1.58 ± 0.53 | 3.65 ± 0.58 | N/A | 4.01 ± 0.69 | N/A |
| **10** | 0.52 ± 0.12 | 3.59 ± 1.27 | 5.76 ± 1.10 | 82.8 ± 40.6 | 42.3 ± 3.53 | 0.116 ± 0.021 |
| **11** | 70.4 ± 31.0 | N/A | N/A | 25.6 ± 21.6 | 81.0 ± 2.77 | 0.004 ± 0.0004 |

We also tested Ac-FLTD-CMK, which is a peptide inhibitor derived from the tetrapeptide sequence of GSDMD, and zVAD-FMK, a broad-spectrum caspase inhibitor (**Figure S12**)^63^. In all cases, zVAD-FMK was the most potent inhibitor, inhibiting all caspases at low – mid nanomolar concentrations (**Figure S12, Table 2**). Ac-LESD-CMK inhibited the extrinsic apoptotic initiator caspases over 100 fold more potently than Ac-FLTD-CMK (**Figure S12B-C, Table 2**). Ac-FLTD-CMK and Ac-LESD-CMK both inhibited caspase-1 at similar IC_50_ values (3.36 μM vs. 5.67 μM, respectively) (**Figure S12D, Table 2**), while both inhibitors weakly inhibited caspase-4, 30 μM vs. 59 μM, respectively, although caspase-4 was difficult to saturate (**Figure S12E, Table 2**). In contrast to the other inflammatory caspases, Ac-LESD-CMK efficiently blocked caspase-5 activity (IC_50_ = 2 μM) better than Ac-FLTD-CMK (IC_50_ = 15 μM) (**Figure S12F, Table 2**). Broadly, mouse caspase-11 and the executioner caspases-3, -6, and -7 were all weakly inhibited by both Ac-FLTD-CMK and Ac-LESD-CMK (**Figure S12G-J, Table 2**). We also tested commercially available caspase-8 inhibitors against caspase-9, and determined that Ac-LESD-CMK was the least potent inhibitor tested (IC_50_ = 12 μM) compared to z-LEHD-FMK (IC_50_ = 1.5 μM), VX-765 (IC_50_ = 4 μM), and z-IETD-FMK (IC_50_ = 3.7 μM). In summary, our new Ac-LESD-CMK tool represents the most selective inhibitor of caspase-8 tested in our assays and thus provides a useful tool compound to explore caspase-8 biology.

### Peptide based caspase inhibitors prevent processing of inflammatory and apoptotic caspase substrates in cells

To determine the activity of the inhibitors in a cellular context, we employed a *Yersinia pseudotuberculosis (Yptb)* model wherein infection leads to caspase-8-mediated processing of caspase-1, which then cleaves and activates GSDMD^54, 57^. We treated *Mlkl* knock-out (KO) mouse bone-marrow derived macrophages (mBMDMs) with 100 µM Ac-LESD-CMK, Ac-FLTD-CMK, or zVAD-FMK for 30 minutes then infected with *Yptb* and tracked the kinetics of cell death by measuring the uptake of Sytox Green dye over time (**Figure 5A**). As anticipated, zVAD-FMK abrogated all cell death, and Ac-LESD-CMK more significantly attenuated cell death compared to Ac-FLTD-CMK (**Figure 5A**), suggesting that Ac-LESD-CMK inhibited caspase-8 activation in cells. Consistent with this, immunoblot analysis demonstrated that Ac-FLTD-CMK failed to prevent caspase-8-mediated activation of caspase-1 and subsequent GSDMD cleavage, but both zVAD-FMK and Ac-LESD-CMK abrogated caspase-1 processing into the mature p20 species and GSDMD processing (**Figure 5B**). Since Ac-LESD-CMK is a potent and selective inhibitor of caspase-8 activity and it can also inhibit caspase-1, this likely explains why it is a better inhibitor than Ac-FLTD-CMK during *Yptb* infection.

**Figure 5.**
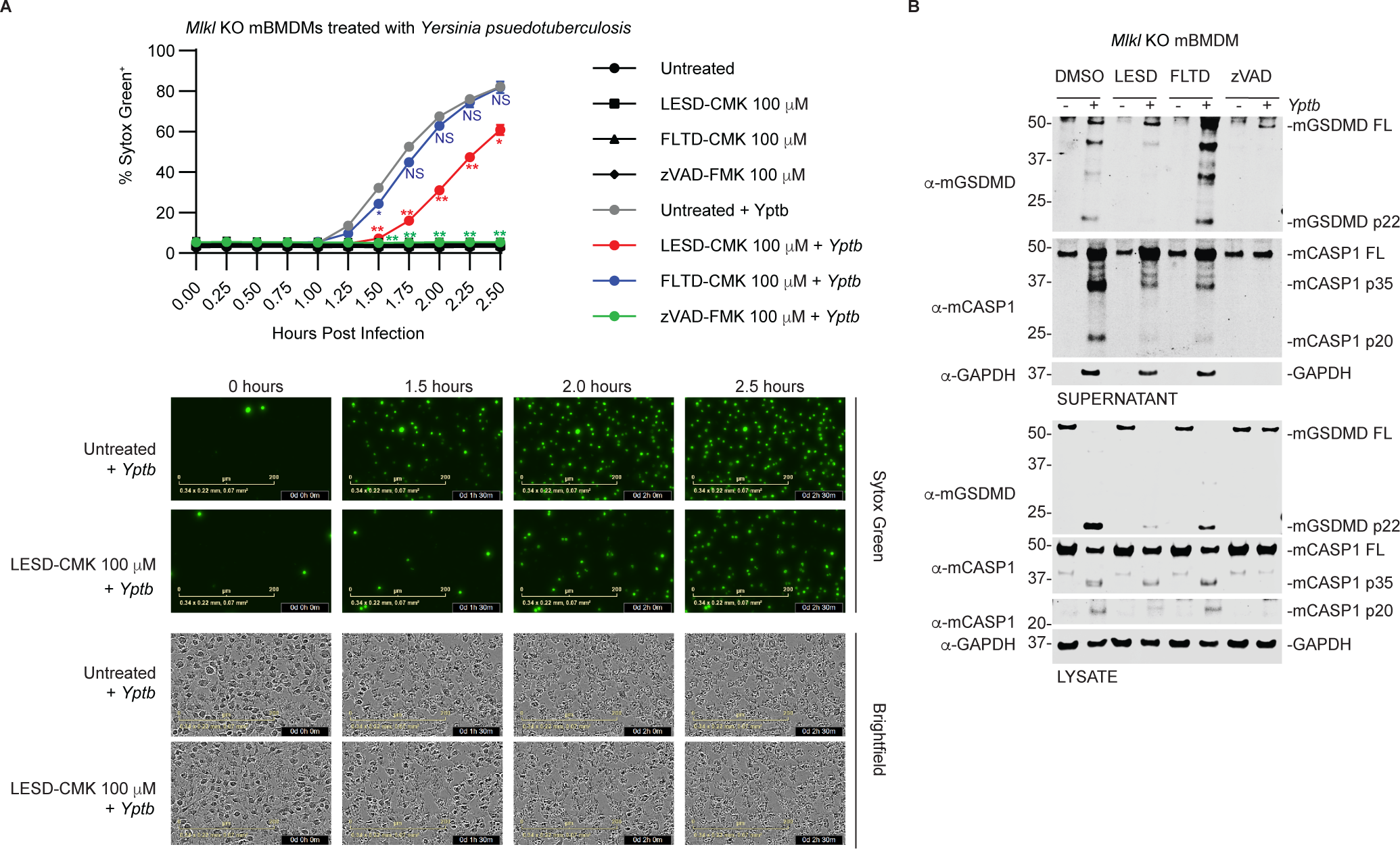
Ac-LESD-CMK inhibits caspase-8 activation in cells. (**A**) Mouse *Mlkl* KO BMDMs were pretreated with 100 μM of the indicated inhibitors for 30 mins before *Yptb* (MOI = 20) infection. The kinetics of cell death was tracked by measuring Sytox Green uptake using an Incucyte. Representative images are depicted below and the quantification is depicted above. (**B**) Immunoblot of *Yptb* infected *Mlkl* KO BMDMs 2.5 h post infection. Data are means ± SEM of three independent experiments. ****P < 0.0001, ***P < 0.001, **P < 0.01, and *P < 0.05 by Two-way ANOVA with Dunnett’s multiple comparison test comparing *Yptb* treatment with no inhibitor to *Yptb* treatment with inhibitors.

To better understand the inhibition profile of Ac-LESD-CMK in cells, we differentiated THP1 monocytes into macrophages and activated the caspase-1 inflammasome using Nigericin (**Figure 6A**), the caspases-4/-5 inflammasomes using LPS (**Figure 6B**), and intrinsic apoptosis using staurosporine (**Figure 6C**)^9, 64^. Before cell death induction, THP1 macrophages were pre-incubated with varying concentrations of Ac-LESD-CMK, Ac-FLTD-CMK, or zVAD-FMK inhibitors for 30 minutes. In agreement with the inhibition potency and pan caspase inhibition profile observed *in vitro,* zVAD-FMK was the most potent dose-dependent inhibitor of pyroptotic cell death upon Nigericin and LPS treatment (**Figure 6A, B**). At all concentrations tested, zVAD-FMK inhibited GSDMD, IL-1β, and caspase-1 activation. zVAD-FMK also inhibited cell death, caspase-3 and PARP cleavage when apoptosis was induced with staurosporine (**Figure 6C**). Consistent with our *in vitro* data, Ac-LESD-CMK did not significantly inhibit apoptosis activated by staurosporine, which activates caspase-9 to induce intrinsic apoptosis. Ac-FLTD-CMK and Ac-LESD-CMK dose-dependently inhibited cell death with LPS treatment (**Figure 6B**) but had a modest impact on cell death with nigericin and staurosporine treatment (**Figure 6A, C**). However, both Ac-FLTD-CMK and Ac-LESD-CMK prevented GSDMD, IL-1β, and caspase-1 activation. Ac-FLTD-CMK appeared to be more potent at inhibiting GSDMD, IL-1β, and caspase-1 processing in cells compared to Ac-LESD-CMK as GSDMD and IL-1β processing was completely abrogated at all concentrations of Ac-FLTD-CMK tested whereas only the highest concentration of Ac-LESD-CMK inhibited GSDMD and caspase-1 processing (**Figure 6A, B**). Similarly, caspase-3 activation, as assessed by the inhibition of caspase-3 processing into the p19 and p17 species, was poorly attenuated with Ac-FLTD-CMK and Ac-LESD-CMK inhibitors, and PARP cleavage was unaffected by either inhibitor (**Figure 6C**). Altogether, our data suggests that Ac-LESD-CMK strongly inhibits activation of caspase-8-dependent cell death and modestly inhibits cell death triggered by inflammasome activation, demonstrating its utility for studying caspase-8 biology in cells.

**Figure 6.**
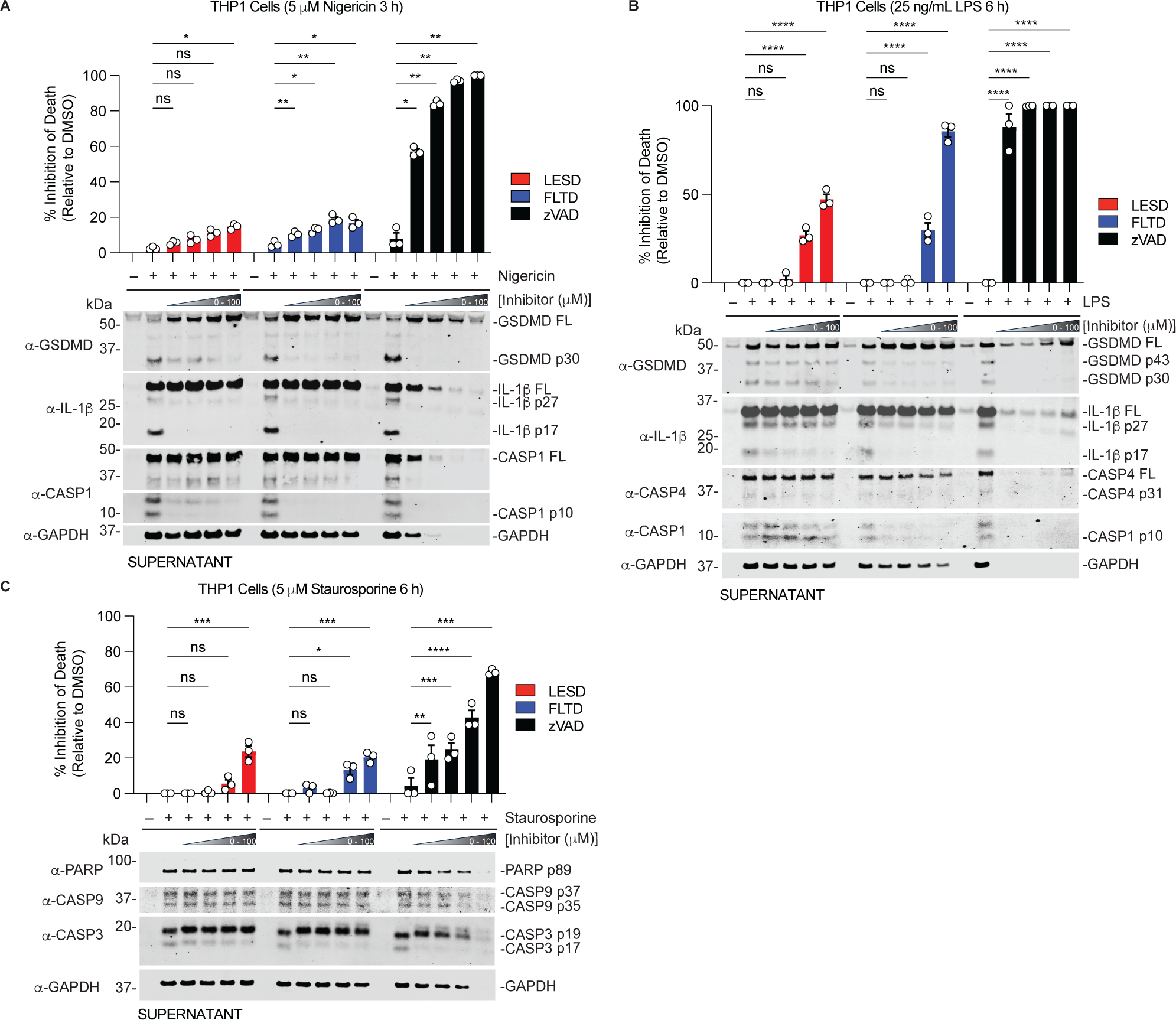
Profiling inhibitors in human macrophages. (**A**) THP1 monocytes were differentiated into macrophages using 50 ng/mL of phorbol 12-myristate 13-acetate (PMA) for two days. Cells were preincubated with the Ac-LESD-CMK, Ac-FLTD-CMK, zVAD-FMK inhibitors at 0, 12.5, 25, 50 or 100 μM, or matched DMSO concentrations for 30 minutes before the addition of 5 μM Nigericin for 3 h. Cell lysis was assessed by Sytox Green uptake. Protein was precipitated from supernatants and analyzed by immunoblotting. (**B**) THP1 cells were treated as in A with inhibitors and non-canonical inflammasome activation triggered by transfection of LPS at 25 ng/mL for 6 h. (**C**) Cells were treated as in A with inhibitors then apoptosis was triggered using 5 μM staurosporine 6 h. Supernatants were precipitated and immunoblotted as in A. Data are means ± SEM of three independent experiments. ****P < 0.0001, ***P < 0.001, **P < 0.01, and *P < 0.05 by two-way ANOVA test.

## Conclusion

The caspases are all structurally similar, making it challenging to develop specific active site inhibitors. Many inhibitors have been reported but at high doses, most have off-target activity for other caspases^41^. Thus, understanding the selectivity and concentrations needed to target certain caspases over others will aid in uncovering their roles in biology. In the present study, we have systematically characterized the relative catalytic efficiency and determined the half-maximal inhibitory concentration of peptide-based substrates and inhibitors of inflammatory and apoptotic caspases. We tested each caspase at the same activity units (as opposed to enzyme concentration) and compared their activities for a given substrate or inhibitor. A limitation of this approach is that activity units are determined using the optimal substrate specific to that caspase, sometimes performed at different temperatures. Without the enzyme concentrations, exact kinetic parameters cannot be measured and while a comparison can be made for different substrates for a given caspase, bias may be introduced when comparing a substrate between caspases. Therefore, we interpret our catalytic efficiencies to be relative.

As anticipated from prior work using positional scanning peptide libraries to determine the specificity of caspases,^33, 36, 37, 65^ Ac-YVAD-AMC, a peptide substrate based on the sequence of IL-1β, was the only substrate that was specifically cleaved by the human inflammatory caspases (caspases-1, -4, -5) but caspases-4 and -5 are unable to cleave this sequence on the native IL-1β substrate^20^, suggesting that additional factors confer specificity for protein substrates. Indeed, it is now known that exosites, alternative binding sites distinct from the active sites, play a critical role in substrate recognition by inflammatory and apoptotic caspases^21, 22, 49-53^. Also in agreement with prior work^37^, the apoptotic caspases also exhibited significantly higher catalytic efficiency for the Ac-DEVD-AMC substrate compared to sequences based on pyroptotic substrates. However, there were differences in the catalytic efficiency amongst the apoptotic caspases for Ac-DEVD-AMC, implying that they may have differences in the ability to process protein substrates.

The observation that substrates and inhibitors derived from inflammatory substrate sequences conferred some selectivity for the inflammatory caspases, and our observation that the LESD sequence in IL-18 played a critical role in IL-18 recognition by the non-canonical inflammasomes (caspases-4/-5), prompted us to develop a specific substrate and inhibitor pair of the inflammatory caspases. Although our new substrate and inhibitor targeted the inflammatory caspases, they were most selective for caspase-8 followed by caspase-10. Notably, this inhibitor did not target caspase-9 and will be a useful tool to distinguish the contributions of extrinsic and intrinsic apoptosis pathways. Caspase-8 is essential for development and plays a role in regulating apoptosis, necroptosis, and has recently been implicated in pyroptosis as well^66, 67^. The existence of a selective caspase-8 inhibitor will facilitate studies into the function of caspase-8 activity in various biological pathways in which caspase-8 is the initiating or dominant protease activated. For example, *Yersinia* infection of macrophages triggers robust activation of caspase-8, which leads to caspase-1 activation due to pathogen blockade of NF-κB signaling^54, 57, 58^. We found that the LESD inhibitor robustly prevented caspase-8 activation and activation of downstream responses. Interestingly, caspase-8 is thought to negatively regulate a constitutively active inflammatory state in neutrophils and its inhibition drives the production of cytokines and chemokines that promote neutrophil mobilization and antibacterial defense *in vivo^68^*, suggesting that caspase-8 inhibition may have therapeutic benefits in certain contexts.

Except for Ac-YVAD-AMC, all other substrates and inhibitors targeted both apoptotic and pyroptotic caspase, and in particular, caspase-8. Even VX-765, which is developed to be a specific caspase-1 inhibitor, inhibited caspase-8, and caspase-8 had high catalytic efficiency against inflammatory peptide substrates such as Ac-WEHD-AMC, and Ac-LEHD-AMC. It is possible that because of the pleotropic effects it has on biology, caspase-8 serves as a backup pathway for the inflammatory caspases and may have overlapping substrate specificities. Consistent with this idea, caspase-8 was demonstrated to form an inflammasome with ASC and NLRC4 when caspase-1 is defective^69^. We did not detect *in vitro* cleavage of IL-1β and IL-18 by caspase-8 but the LESD-based tetrapeptide substrate and inhibitor targets caspase-8. Other factors such as exosites, subcellular localization, and presence of pathogenic factors may also influence the substrate repertoire and activity of caspases in cells.

It was somewhat surprising that the LESD-based inhibitor was selective for casapase-8. Several factors can account for this, such as conformational changes upon substrate binding, sterics, and hydrophobic and electrophilic interactions at the active site between the peptide substrate and residues in the caspase-8 active site. For example, mouse caspase-11 lacks key electrophilic interactions with the LESD sequence, and harbors an elongated loop, compared to the human caspase-4 ortholog, that sterically occludes IL-18 and prevents its processing^22^. Prior work on caspase-9 suggests that in addition to the sequence, the local context and three-dimensional positioning and interactions influences substrate processing^70^. Some sequences can be recognized but are not processed unless the correct local environment is present. This might explain why other caspases such as caspases-4/-5 and -6 are unable to process the YVHD sequence in IL-1β but are able to process IL-1β when the YVHD sequence is substituted with LESD. Structural studies are needed to better understand the basis of the selectivity of LESD for caspase-8.

Currently, z-IETD-FMK is thought to be a selective caspase-8 inhibitor^68^. However, we observed that Ac-LEHD-FMK was more potent than z-IETD-FMK at inhibiting caspase-8 activity, in line with a prior report that caspase-8 processed an LEHD-based probe 4.5 fold faster than the corresponding IETD probe^39^. However, the assumption that substrate selectivity should match inhibitor selectivity is inaccurate as our data indicates that caspase-6 is inhibited by an LESD-based inhibitor despite being unable to cleave the Ac-LESD-AMC substrate. Ac-LEHD-FMK was not selective and exhibited potent inhibition of other caspases, such as caspases-1, -3 and - 7. Notably, our LESD-based inhibitor was also more potent at inhibiting caspase-8 compared to z-IETD-FMK and was not as potent at inhibiting other caspases like Ac-LEHD-FMK, indicating that it is both potent and selective for caspase-8. Although Ac-LESD-AMC was exceptionally potent *in vitro*, it appeared to be less potent in cells as high doses were needed to see cellular inhibition. This is likely due to a lack of cell penetrance and efforts are underway to synthesize a more cell penetrant version to facilitate biological studies.

A peptide inhibitor based on the sequence of GSDMD, Ac-FLTD-CMK, was reported to be specific for inflammatory caspases^63^. We observed a similar trend in inhibition profile but the IC_50_ values we determined were higher in magnitude. For example, Yang *et* al. reported an IC_50_ value of 0.0467 µM for caspase-1^63^ whereas we measured an IC_50_ value of 3.38 µM. This is likely because they used 50 nM of enzyme and 5 µM of the Ac-WEHD-AMC substrate for activity when measuring the IC_50_ value whereas we used 0.25 activity units of caspase-1 and 200 µM Ac-WEHD-AMC substrate in our study. We also discovered that the Ac-FLTD-CMK also inhibits caspase-8 but this inhibition is 100-fold weaker than our Ac-LESD-CMK inhibitor. Overall, our findings support the known selectivity of peptide substrates and inhibitors and offers some new insights, such as the finding that VX-765 inhibits caspase-8, which was not previously known. We anticipate that this database of substrate and inhibitor selectivity for caspases assayed at the same activity units, along with our new selective probes for caspase-8, will prove useful for selecting pharmacological tools to facilitate future studies into the function of caspases in biology.

## Method Details

### Antibodies and reagents table

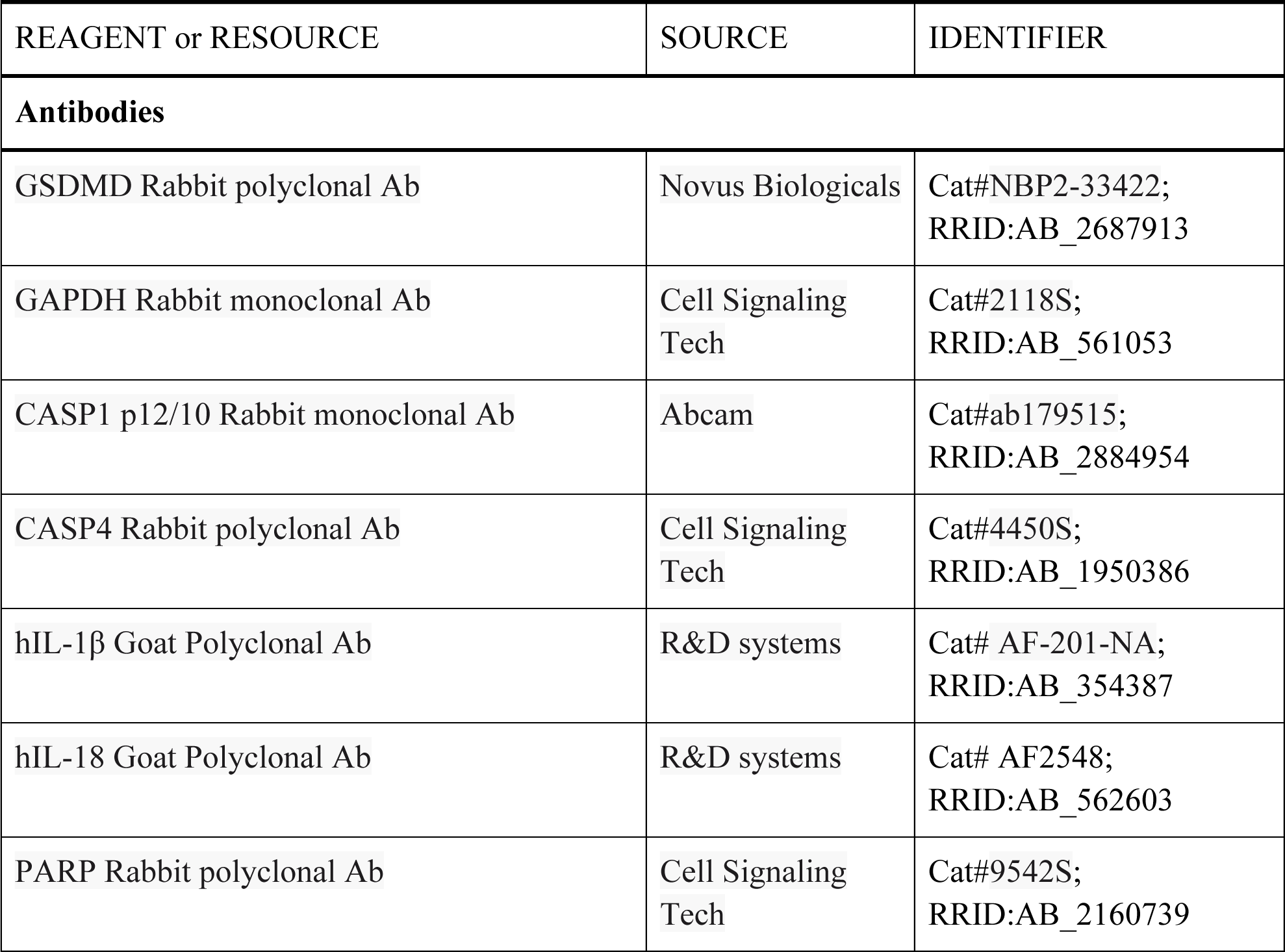
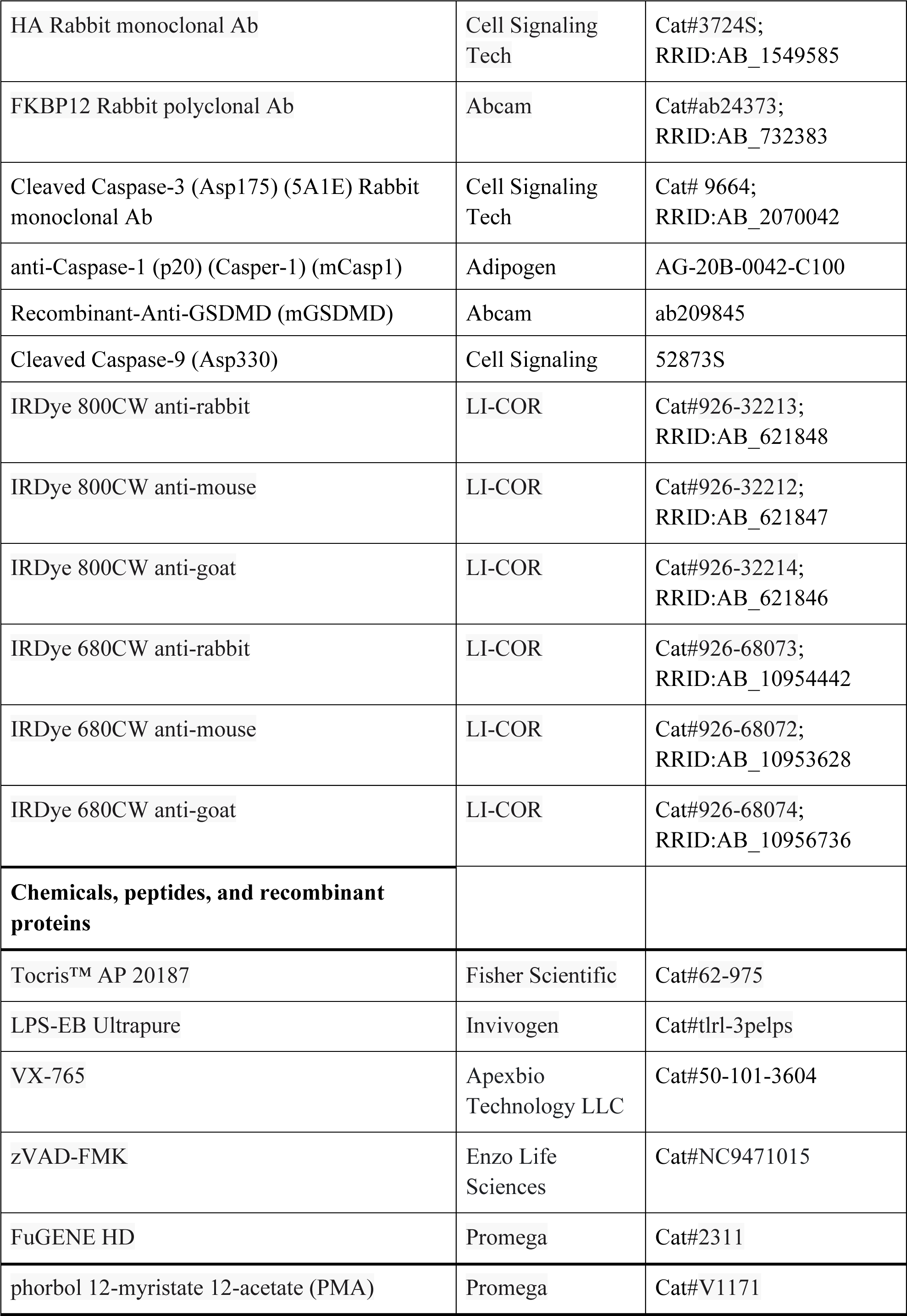
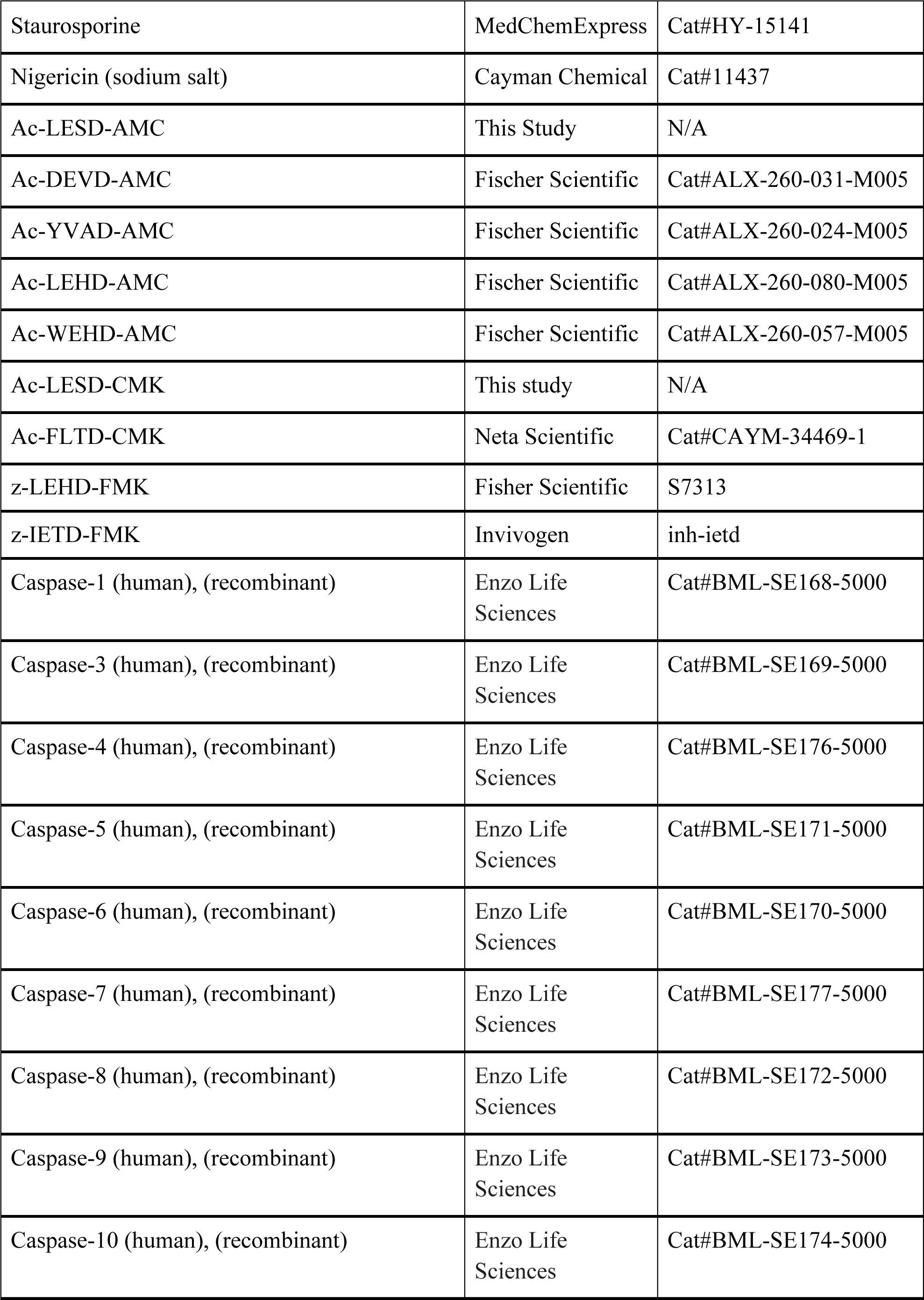
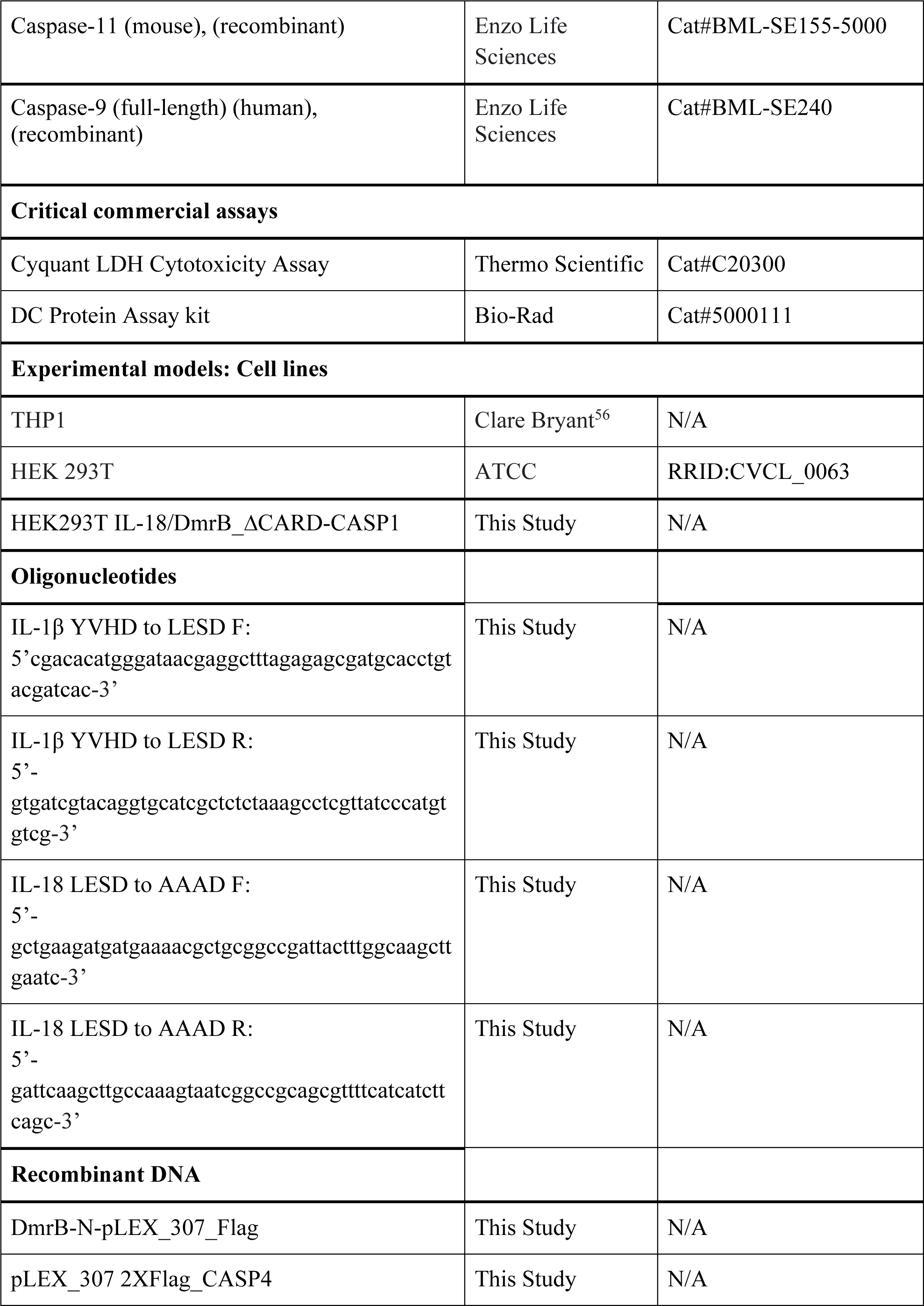
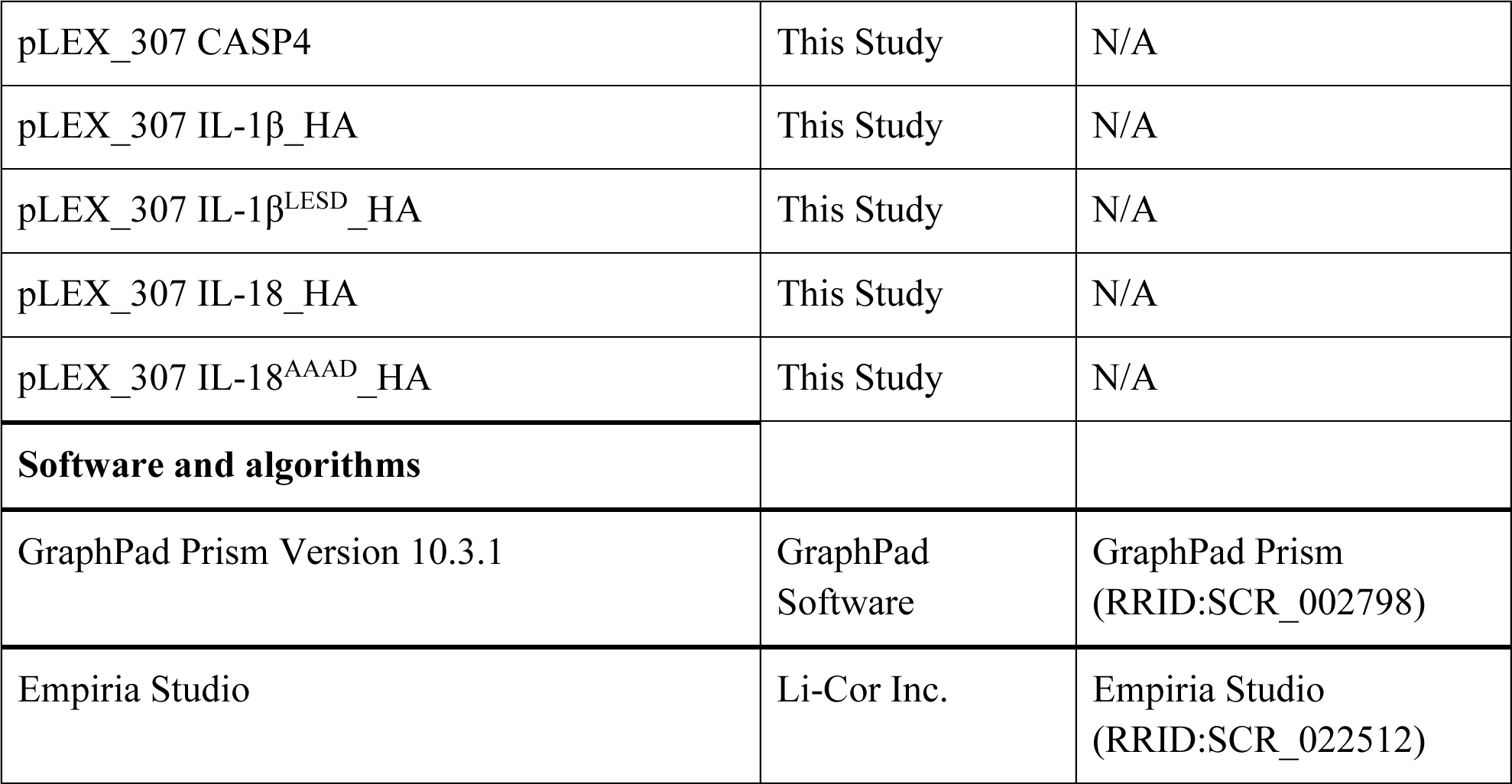

### Recombinant Caspase Assays

All *in vitro* assays using recombinant caspases were performed in 20 μL reactions containing caspase assay buffer (20 mM PIPES, 100 mM NaCl, 10 mM DTT, 1mM EDTA, 0.1% CHAPS, 10% sucrose, pH 7.). We used 0.25 activity units/μL of each indicated caspase. For cytokine cleavage assays, 10 ug of each were transfected into 10 cm plates of HEK 293T cells using FuGENE HD Transfection Reagent (Promega, E2312) according to the manufacturer’s protocol. After 24 hours, cells were harvested and lysed by sonication for 10 second pulses for 30 seconds at 30% amplitude in PBS and centrifuged at 12,000 g to remove cell debris. Lysates were diluted 10x in caspase assay buffer, and aliquoted for incubation with caspases or further purified as previously described^20^ before cleavage assays. At the indicated time points, samples were mixed 1:1 with 2x Licor Protein Sample Loading Buffer (Neta Scientific, 928-40004) and boiled at 95 °C for 10 minutes, then analyzed by SDS-PAGE and immunoblotting.

#### In Vitro Substrate Kinetics

All fluorogenic substrates were resuspended in DMSO at 40 mm stock concentrations and diluted to the indicated concentrations in caspase assay buffer. All probe experiments were completed at 20 µL final volume, with 0.25 activity units per µL of each recombinant caspase (1 activity units per µL for caspase-9). Relative fluorescence units were measured every 5 minutes using a Cytation 5 plate reader (Biotek). Background fluorescence was subtracted from samples incubated with caspases. At each concentration of substrate, simple linear regressions (y = mx + b) using GraphPad Prism were performed to determine the rate. Rates were plotted against substrate concentration for n = 3 separate runs and fit to a Michaelis-Menten curve (Y = V_max_*X/(*K*_M_ + X)) on Prism. Caspase-substrate pairs that resulted in V_max_ values under 20 RFUs/minute were considered undetected. For V_max_, *K*_M_ and *k*_cat_/*K*_M_ calculations, curves were generated for each individual experimental replicate, and V_max_, *K*_M_, and *k*_cat_ values were recorded for each replicate. *k*_cat_ was determined as the V_max_ value with total enzyme concentration (ET) set to 1 for all caspases. The mean and standard error mean were calculated by compiling the values for each individual run.

#### In vitro Inhibition Assays

To determine inhibition of maximal enzyme activity, each caspase was incubated with its optimal probe substrate at a saturating concentration. Inhibition of maximal caspase activity was determined with 200 µM Ac-WEHD-AMC for caspase-1 and -5, 1000 µM Ac-WEHD-AMC for caspase-4 and -11, Ac-LEHD-AMC at 200 µM for caspase-8 and -10, Ac-DEVD-AMC at 25 µM for caspase-3 and -7 with ZVAD-FMK, and Ac-DEVD-AMC at 100 µM for caspase -3, -6, and -7 for all other inhibitors tested, and 200 µM Ac-LEHD-AMC for caspase-9. Rates were determined as described in the *In Vitro Substrate Kinetics* section. Rates for n = 3 independent runs were plotted against inhibitor concentration. Percent caspase activity was determined by normalizing each independent run with 0 % activity set at 0, and maximum activity set as the sample with no inhibitor added. Replicates were combined and inhibition curves were calculated for the aggregate data using a [Inhibitor] vs. normalized response curve on Graphpad Prism (Y=100/(1+X/IC_50_)).

### Testing Inhibitors in Cells

THP-1 cells were resuspended in RPMI medium containing 50 ng/mL phorbol 12-myristate 12-acetate (PMA). Cells were plated in 96-well plates at a density of 8 x 10^4 cells/well. After 48 hours, the media was then replaced with 100 µL per well of Opti-MEM (Thermo Fisher Scientific, 31-985-062) containing Sytox Green (Fisher Scientific, S7020) at 200 nM. Where indicated, cells were treated with Ac-LESD-CMK, Ac-FLTD-CMK or zVAD-FMK, 30 min before the cell death trigger was added (described below). Cells were incubated in the trigger for indicated time point and Sytox green values were read using the Cytation 5 plate reader. Supernatants were harvested and pooled across replicates and proteins were precipitated. Nigericin (Neta Scientific, 11437) was resuspended in ethanol and added at a final concentration of 5 µM while Staurosporine (MedChem Express, HY-15141) was resuspended in DMSO and added at a final concentration of 1 µM. The LPS transfection solution was prepared by adding LPS (Cell Signaling Technology, 14011S) (25 mg/mL final concentration) and FuGENE (Promega, E2312) (0.5% final concentration) to Opti-MEM. This solution was gently mixed by flicking and incubated for 15 min at room temperature before addition to each well.

### Western Blotting

Protein samples were run on Nupage 4-12% Bis-Tris Midi Protein Gel (Thermo Scientific, WG1403BOX) at 175 volts. Proteins were transferred to 0.45 µm nitrocellulose membranes (BioRad,1704271) at 25 volts for 7 minutes on the Biorad Transblot Turbo System. Membranes were blocked using LICOR Intercept blocking buffer (Neta scientific, 927-70010) for 1 hour and stained with primary antibodies at 1:1000 concentration in 50% Licor blocking buffer and 50% TBS buffer with 0.1% Tween for 1 hour at room temperature or overnight at 4C. Membranes were washed 3x 15 minutes, followed by incubation with Donkey Anti-Mouse/Rabbit/Goat IgG Polyclonal Antibody (IRDye® 800CW) for 1 hour at room temperature. Membranes were then washed 3x10 minutes and imaged on an Odyssey M Imaging System (LI-COR Biosciences). Images were analyzed with Empiria studio version 3.2 (Licor), and brightness, contrast, and tone parameters were adjusted uniformly for entire membranes on Adobe Photoshop.

### Cell Culture

HEK 293T and THP1 cells were purchased from ATCC. HEK 293T cells were cultured in Dulbecco’s Modified Eagle’s Medium (DMEM) with L-glutamine and 10% fetal bovine serum (FBS). THP-1 cells were cultured in Roswell Park Memorial Institute (RPMI) medium 1640 with L-glutamine and 10% fetal bovine serum (FBS). Isolated bone marrow cells from 6- to 10-week-old male and female mice were grown at 37°C and 5% CO2 in 30% macrophage medium [30% L929 fibroblast supernatant and complete Dulbecco’s modified Eagle’s medium (DMEM)]. BMDMs were harvested in cold phosphate-buffered saline (PBS) on day 7 and replated in 10% macrophage medium onto tissue culture (TC)–treated plates or glass coverslips in TC-treated plates. All cells were grown at 37 °C in a 5% CO_2_ atmosphere incubator.

### Generation of stable cell lines

For generating HEK 293T cells ectopically expressing DmrB-caspase 8 and IL-18 or DmrB-caspase-9, plasmids encoding those proteins were packaged into lentivirus by transfecting the vectors (2 µg) along with psPAX2 (2 µg), and pMD2.G (1 µg) using Fugene HD transfection reagent (Promega) into HEK 293T cells. After 2 days, the supernatants were filtered using a 0.45 µm filter, then used to infect HEK 293T cells. After 48 hours, the cells expressing the indicated constructs were selected with blasticidin (10 µg/mL), or puromycin (1 µg/mL). During transfection and transduction of DmrB-caspase-8 into the HEK 293T IL-18 cell line, 40 μM of zVAD-FMK was incubated with the cells to prevent spurious apoptosis to promote sufficient production and integration of the virus.

### Cloning

All plasmids were cloned using Gateway technology as previously described^20^. DNA encoding the indicated proteins were inserted between the *attR* recombination sites and shuttled into modified pLEX_307 vectors (Addgene) using Gateway technology (Thermo Fisher Scientific) according to the manufacturer’s instructions. Proteins expressed from these modified vectors contain an N-terminal *att*B1 linker (GSTSLYKKAGFAT) after any N-terminal tag (e.g. DmrB) or a C-terminal *att*B2 linker (DPAFLYKVVDI) preceding any C-terminal tag such as V5 or HA. ΔDED-1/2 DmrB-caspase-8 beginning at residue 146 (M146) or ΔCARD DmrB-caspase-9 beginning at residue 139 (V139), which was changed from valine to methionine, were cloned into a modified pLEX_307 vector and contained the N-terminal *att*B1 linker between the DmrB and caspase sequences.

### *Yersinia pseudotuberculosis* primary murine bone marrow-derived macrophage infections

*Yersinia pseudotuberculosis (Yptb)* infections were performed as previously described^54^. Strain IP2666 was induced before infection by diluting stationary phase overnight cultures 1:40 in 3 ml of inducing medium[2× Yeast Extract Tryptone (YT) broth, 20 mM sodium oxalate, and 20 mM MgCl2]. Cultures were grown at 28°C for 1 hour and transferred to 37°C for 2 hours shaking. Bacterial growth was measured by absorbance at optical density at 600 nm (OD600) using a spectrophotometer. Bacteria were pelleted, washed, and resuspended in Opti-MEM or 10% macrophage media (10% L929 fibroblast supernatant and complete Dulbecco’s modified Eagle’s medium (DMEM)) for infection. *In vitro* infections were performed at multiplicity of infection of 20. Indicated inhibitors were added 30 minutes prior to bacterial addition and plate was incubated at 37°C. Gentamycin (100 μg/ml) was added 1 hour after infection for all infections. Plate was imaged in a Sartorius Incucyte S3.

## Synthesis of Ac-LESD-AMC substrate and Ac-LESD-CMK inhibitor

### General Peptide Synthesis

Solvents and reagents were purchased from commercial suppliers and used without further purification. Analytical LC-MS was performed using a system comprised of an Agilent 1260 Infinity II HPLC instrument equipped with an Agilent InfinityLab LS/MSD XT MS detector with electrospray ionisation. The system ran with an Agilent ZORBAX SB-C18 RRHT (50 mm × 4.6 mm × 1.8 μm) column and gradient elution with two binary solvent systems: MeCN/H_2_O or MeCN/H_2_O plus 0.1% formic acid. Preparative HPLC was performed using a system comprising an Agilent 1260 Infinity II HPLC system equipped with an Agilent 1290 Prep Bin Pump, an Agilent Prep-C18 column (250 mm × 21.2 mm × 10 μm), and an Agilent prep autosampler and fraction collector. The system ran using a UV diode array detector and purification was performed using a gradient elution using a MeCN/H_2_O binary solvent system containing 0.05% trifluoroacetic acid.

### Ac-LESD-AMC

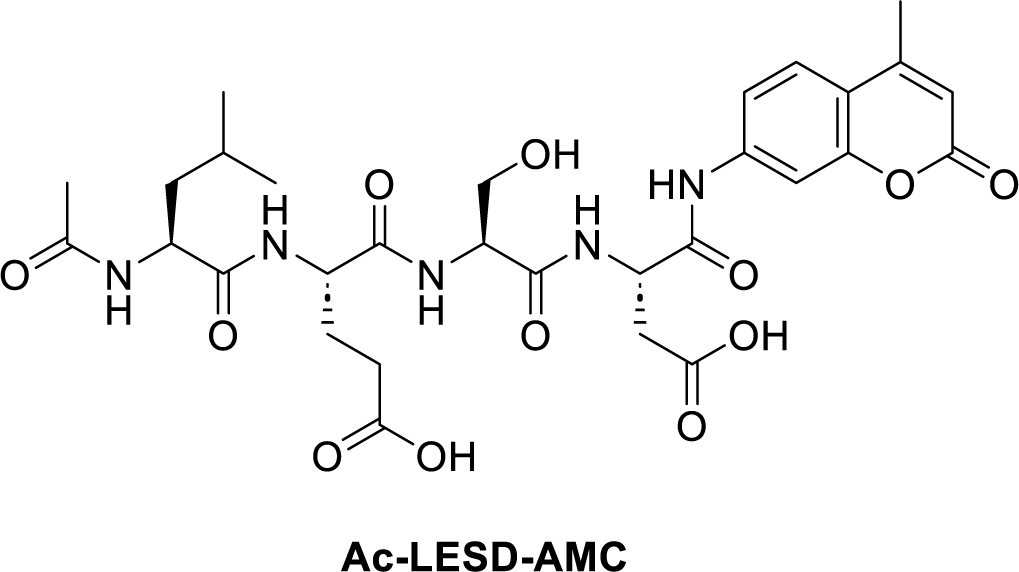

**Ac-LESD-AMC** was synthesized by manual Fmoc solid phase peptide chemistry using Fmoc-Asp(Wang-resin)-AMC. Following resin swelling in DMF, iterative cycles of Fmoc deprotection (20% piperidine in DMF containing 0.1 M HOBt, 3 x 5 minutes), amino acid coupling (4 eq. Fmoc AA, 4 eq. HATU in 4% DIPEA in DMF, 1 hour) and washing (3 x DMF, 3 x DCM) were performed until the sequence was complete. N-terminal acetylation was achieved using 5% acetic anhydride in pyridine. Peptides were deprotected and cleaved from resin by treatment with trifluoroacetic acid/water/triisoproylsilane/phenol (88:5:5:2) for 1-3 hours. Solvent was removed under a flow of nitrogen and peptides were precipitated with ice cold ether. **Ac-LESD-AMC** was subsequently purified by reverse phase chromatography eluting with 5-95% acetonitrile in water containing 0.05% trifluoroacetic acid over a C18 column. Peaks containing **Ac-LESD-AMC** were identified by LC-MS, pooled, and concentrated *in vacuo*.

Ac-LESD-AMC: *m*/*z* [M + H]^+^ calcd for C_30_H_39_N_5_O_12_ 661.3; found 661.9

### Ac-LESD-CMK

**Scheme 1.**
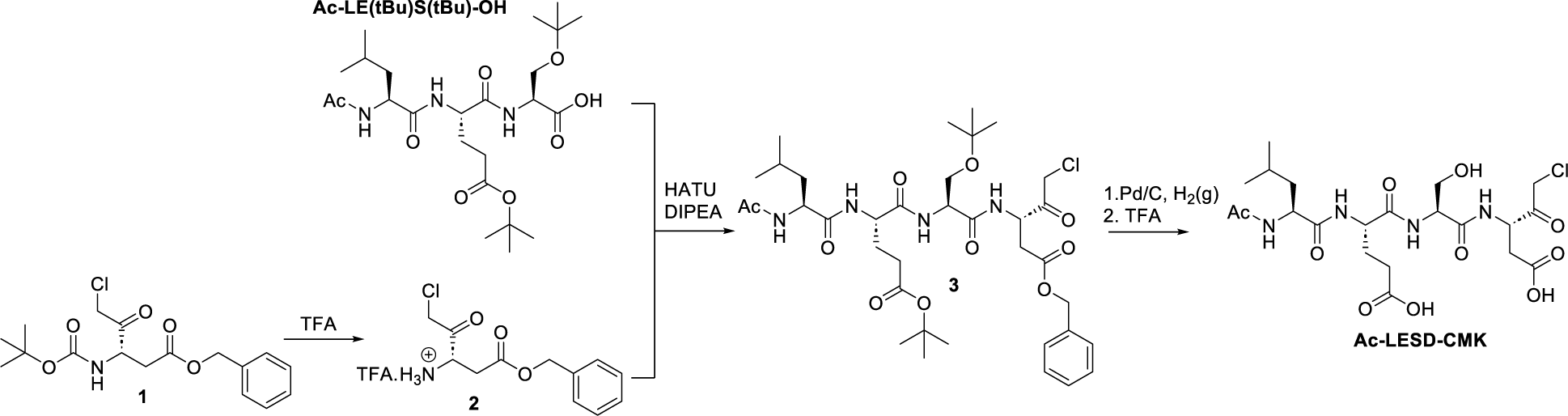
Synthesis of Ac-LESD-CMK.

**Ac-LE(tBu)S(tBu)-OH** was synthesized by manual Fmoc solid phase peptide chemistry on 2-chlorotrityl resin. Following resin swelling in DMF, Fmoc-L-Ser(tBu)-OH was loaded onto the resin (1.2 eq AA, 2.5 eq DIPEA in DCM for 1 hour), the resin was capped (3 eq. MeOH, 2.5 eq DIPEA in DCM for 1 hour). The resin then underwent iterative cycles of Fmoc deprotection (20% piperidine in DMF containing 0.1 M HOBt, 3 x 5 minutes), amino acid coupling (4 eq. Fmoc AA, 4 eq. HATU in 4% DIPEA in DMF, 1 hour) and washing (3 x DMF, 3 x DCM) until the sequence was complete. N-terminal acetylation was achieved using 5% acetic anhydride in pyridine. **Ac-LE(tBu)S(tBu)-OH** was cleaved from the resin in 20% hexafluoroisopropanol (HFIP) in DCM at room temperature for 1 hour. Solvent was removed under a flow of nitrogen and the crude material used with no further purification.

Separately, Boc-L-aspartic acid β-benzyl ester chloromethylketone **1** (65 mg, 0.18 mmol) (purchased from Santa Cruz Biotechnologies, Cat # sc-285170) was dissolved in 3 mL DCM at room temperature before the addition of 1 mL trifluoroacetic acid (TFA) and stirred for 1 hour. The solvent and volatiles were removed under reduced pressure. Crude deprotected L-aspartic acid β-benzyl ester chloromethyl ketone **2** was dissolved in 3 mL of tetrahydrofuran (THF) with HATU (68 mg, 1.1 eq) and DIPEA (440 uL, 5.5 eq). To this solution, Ac-LE(tBu)S(tBu)-OH peptide (80 mg, 1 eq) in 1 mL of THF was added. The reaction stirred at r.t. for 2 hours and reaction progress monitored by LC/MS. Solvent and volatiles were removed under reduced pressure and product was redissolved in 10 mL ethyl acetate before 2 x 10 mL washes of 0.5 M HCl. Organic extracts were dried over magnesium sulfate, and concentrated *in vacuo* to yield **3,** which was used with no further purification in the next step.

To a solution of **3** (127 mg, 1 eq.) in THF (10 mL) under an argon atmosphere was added 10% Pd/C (13 mg, 10% w/w) and hydrogen gas was bubbled through the reaction for 2 hours. The reaction mixture was filtered over celite, and the filter cake washed with THF and MeOH. Solvents and volatiles were removed under reduced pressure and dissolved directly in 50% TFA in DCM and stirred at room temperature for 1 hour to generate **Ac-LESD-CMK**. Solvents and volatiles were removed under reduced pressure. **Ac-LESD-CMK** was subsequently purified by reverse phase chromatography eluting with 5-95% acetonitrile in water containing 0.05% trifluoroacetic acid over a C18 column. Peaks containing **Ac-LESD-CMK** were identified by LC-MS, pooled, and concentrated *in vacuo*.

Ac-LESD-CMK: *m*/*z* [M + H]^+^ calcd for C_21_H_33_ClN_4_O_10_ 537.2; found 537.2

## Author contributions CRediT

Conceptualization: CYT, Methodology: CYT, GMB, CMB, NRR, WY, IEB, Investigation: CMB, NRR, MK, PME, SL, WY, TW, RP, MK, TS, MK, BMD, Visualization: CYT, CMB, Funding acquisition: CYT, GMB, IEB, CMB, PME, NRR, Project administration: CYT, GMB, CMB, Data curation: CYT, CMB, Formal Analysis: CYT, CMB, TW, Visualization: CYT, CMB, Resources: CYT, GMB, IEB, Supervision: CYT, GMB, IEB, Writing – original draft: CYT, CMB, Writing – review & editing: CYT, IEB, GMB, CMB, MK, NRR, PME.

## Acknowledgments

We thank all the members of the Taabazuing lab for scientific discussion. This work was supported by an NIH R00 Career Transition Award Grant# 4R00AI148598-03 (CYT), NIGMS Maximizing Investigator’s Research Award (MIRA) Grant# 1R35GM155239-01 (CYT) and Center for Translational Chemical Biology Pilot Award from the University of Pennsylvania (CYT, GMB). GMB is also supported by the National Institutes of Health R35-GM142505. NRR is supported by an NIH Chemistry Biology Interface Training Grant (T32 GM133398). PME is supported by the Martin and Pamela Winter Infectious Disease Fellowship. CMB is a Penn Provost Postdoctoral Fellow and is supported by Burroughs Wellcome Fund (Grant# 1054907). IEB is supported by two NIH R01 grants (Grant# R01AI128530, R01139102A1) and the Mark Foundation.

