## Supplemental Figures for "Chemical Tools Based on the Tetrapeptide Sequence of IL-18 Reveals Shared Specificities between Inflammatory and Apoptotic Initiator Caspases"

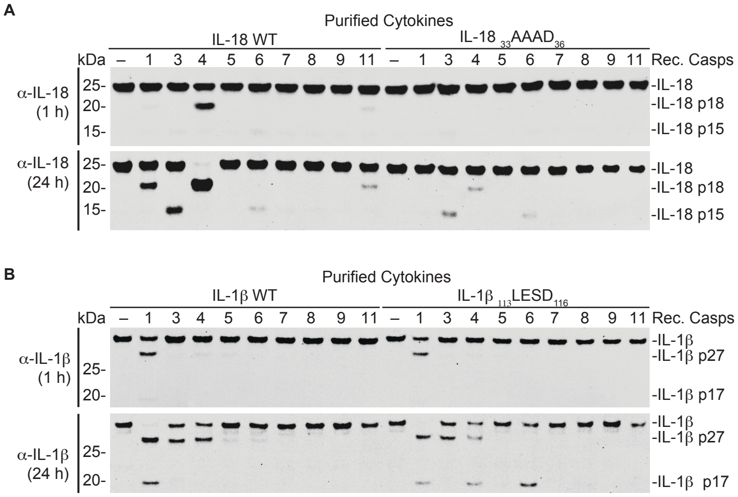


**Figure S1 (Related to Fig. 1). Recombinant caspases cleave purified IL-18 and IL-1β in a tetrapeptide sequence-dependent manner**. (**A**). HA-tagged IL-18 WT and IL-18 with the tetrapeptide sequence at position 33-36 mutated to AAAD (IL-18 AAAD) were expressed and purified from HEK 293T cells, mixed with 0.25 activity units/μL of indicated caspases for 1 or 24 hours before immunoblot analysis. (**B**) IL-1β WT and IL-1β with the tetrapeptide sequence at position 113-116 mutated to LESD (IL-1β LESD), were expressed and purified from HEK 293T cells and treated as in A. Data are representative of three or more independent experiments.

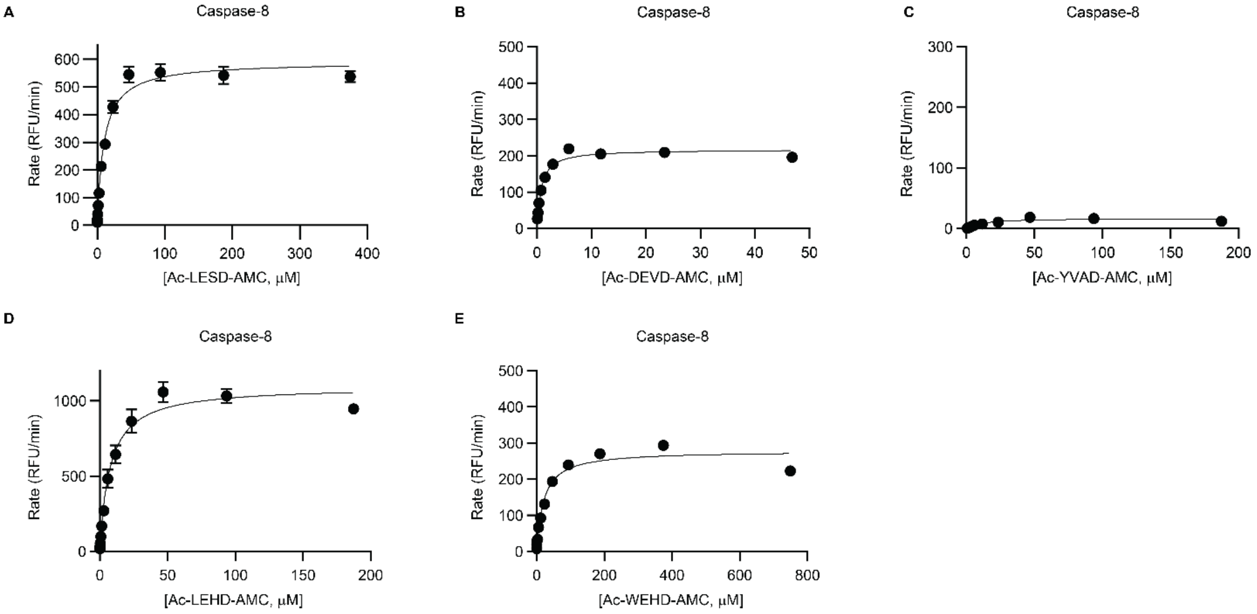


**Figure S2 (Related to Fig. 2). Kinetic characterization of the extrinsic apoptotic initiator caspase-8 with tetrapeptide probes.** (**A-E**) Michaelis-Menten kinetic profiles of (**A**) Ac-LESD-AMC, (**B**) Ac-DEVD-AMC, (**C**) Ac-YVAD-AMC, (**D**) Ac-LEHD-AMC, and (**E**) Ac-WEHD-AMC cleavage by 0.25 activity units/μL of recombinant caspase-8. Data was fitted to the Michaelis-Menten equation in GraphPad Prism and are means ± SEM of three independent experiments.

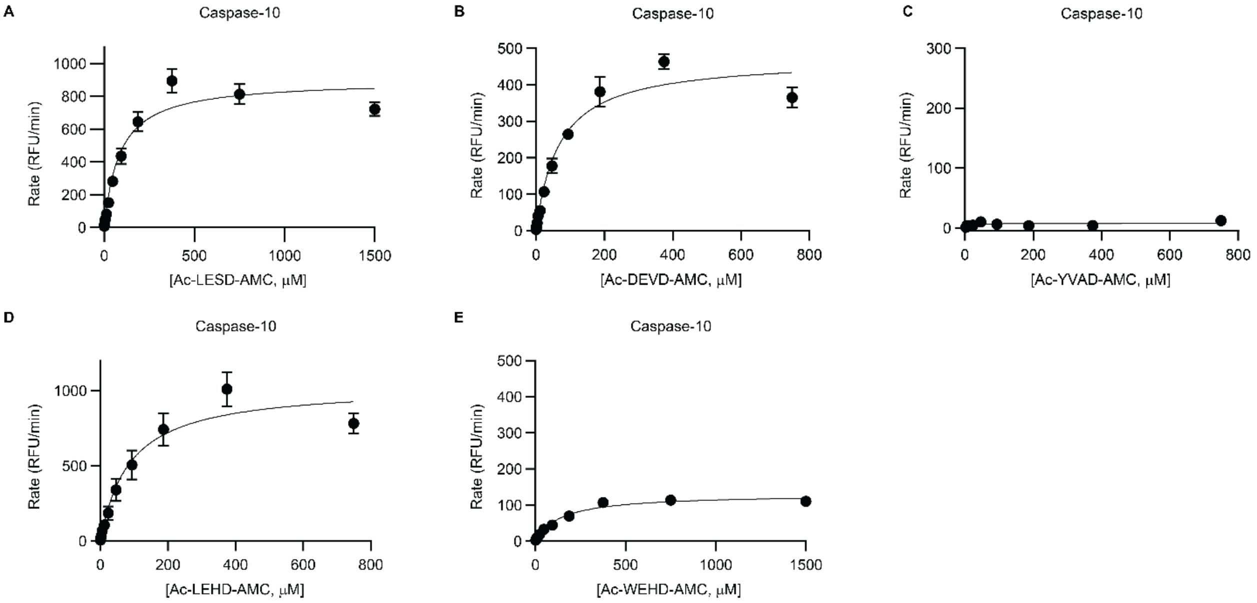


**Figure S3 (Related to Fig. 2). Kinetic characterization of the extrinsic apoptotic initiator caspase-10 with tetrapeptide probes.** (**A-E**) Michaelis-Menten kinetic profiles of (**A**) Ac-LESD-AMC, (**B**) Ac-DEVD-AMC, (**C**) Ac-YVAD-AMC, (**D**) Ac-LEHD-AMC, and (**E**) Ac-WEHD-AMC cleavage by 0.25 activity units/μL of recombinant caspase-10. Data was fitted to the Michaelis-Menten equation in GraphPad Prism and are means ± SEM of three independent experiments.

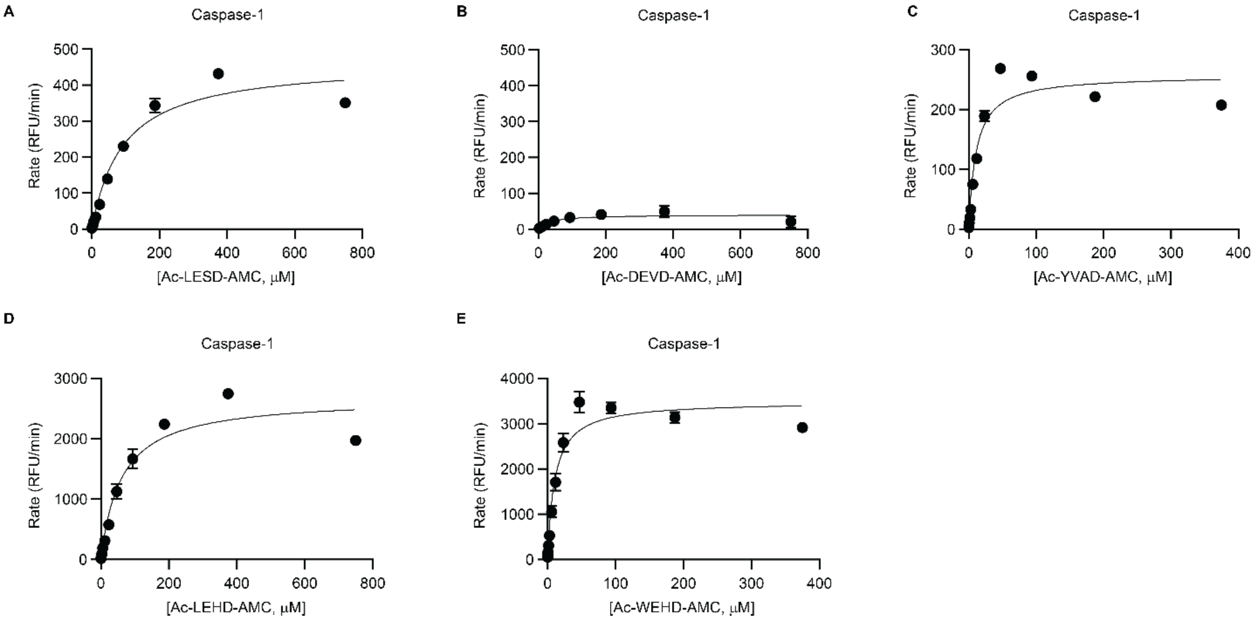


**Figure S4 (Related to Fig. 2). Kinetic characterization of the inflammatory caspase-1 with tetrapeptide probes.** (**A-E**) Michaelis-Menten kinetic profiles of (**A**) Ac-LESD-AMC, (**B**) Ac-DEVD-AMC, (**C**) Ac-YVAD-AMC, (**D**) Ac-LEHD-AMC, and (**E**) Ac-WEHD-AMC cleavage by 0.25 activity units/μL of recombinant caspase-1. Data was fitted to the Michaelis-Menten equation in GraphPad Prism and are means ± SEM of three independent experiments.

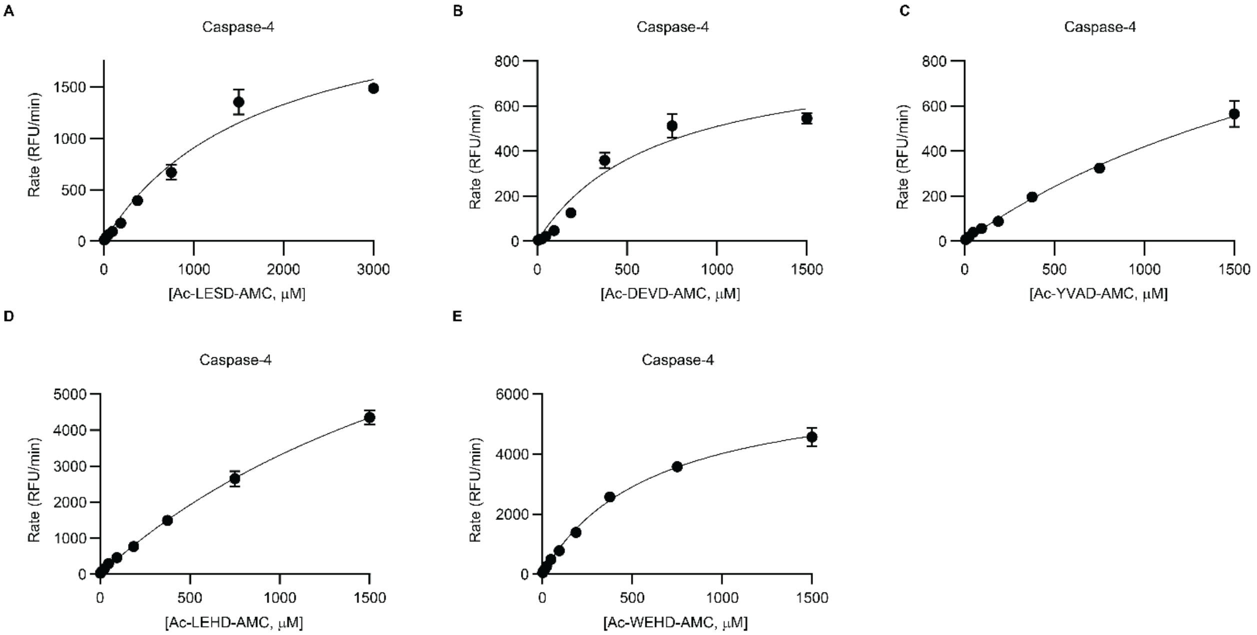


**Figure S5 (Related to Fig. 2). Kinetic characterization of the inflammatory caspase-4 with tetrapeptide probes.** (**A-E**) Michaelis-Menten kinetic profiles of (**A**) Ac-LESD-AMC, (**B**) Ac-DEVD-AMC, (**C**) Ac-YVAD-AMC, (**D**) Ac-LEHD-AMC, and (**E**) Ac-WEHD-AMC cleavage by 0.25 activity units/μL of recombinant caspase-4. Data was fitted to the Michaelis-Menten equation in GraphPad Prism and are means ± SEM of three independent experiments.

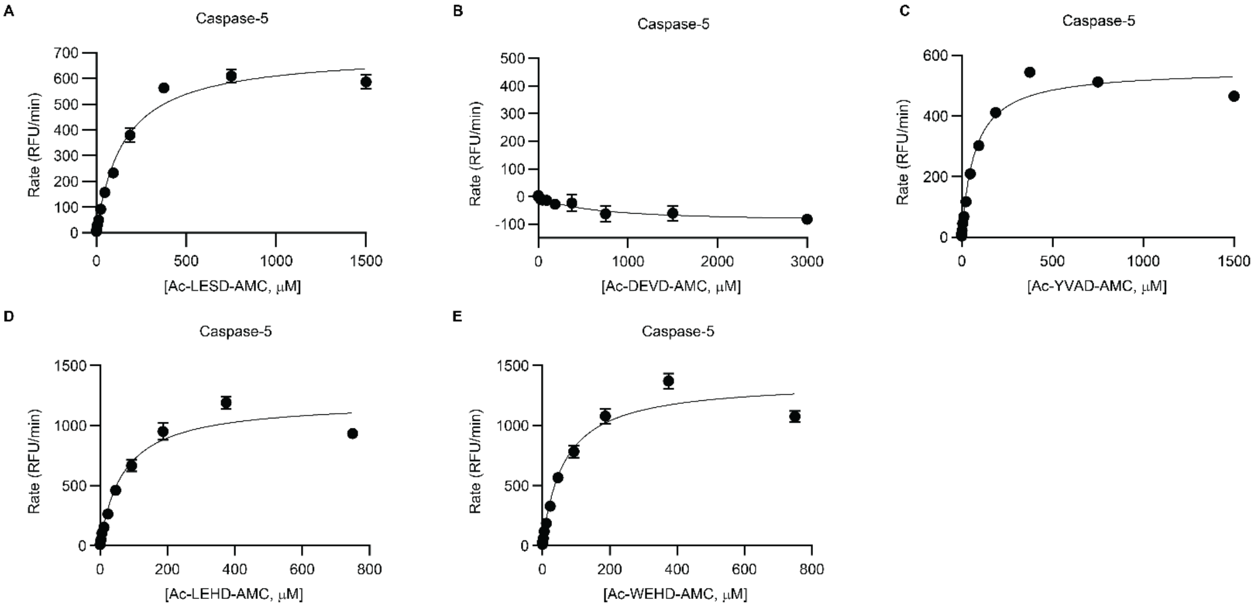


**Figure S6 (Related to Fig. 2). Kinetic characterization of the inflammatory caspase-5 with tetrapeptide probes.** (**A-E**) Michaelis-Menten kinetic profiles of (**A**) Ac-LESD-AMC, (**B**) Ac-DEVD-AMC, (**C**) Ac-YVAD-AMC, (**D**) Ac-LEHD-AMC, and (**E**) Ac-WEHD-AMC cleavage by 0.25 activity units/μL of recombinant caspase-5. Data was fitted to the Michaelis-Menten equation in GraphPad Prism and are means ± SEM of three independent experiments.

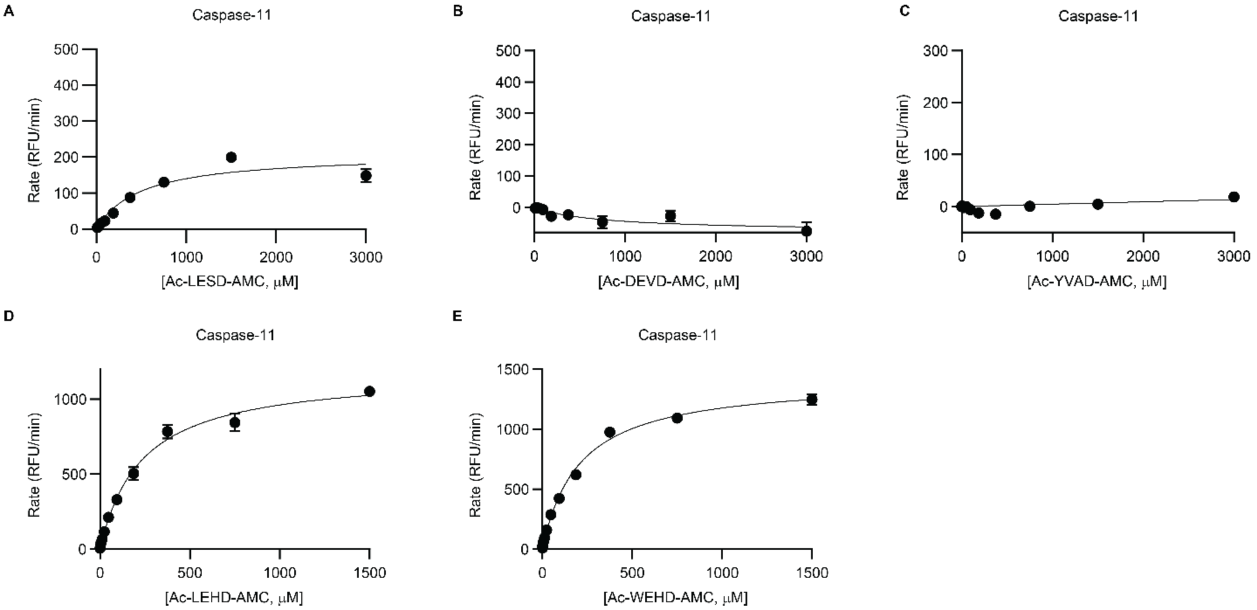


**Figure S7 (Related to Fig. 2). Kinetic characterization of the inflammatory mouse caspase-11 with tetrapeptide probes.** (**A-E**) Michaelis-Menten kinetic profiles of (**A**) Ac-LESD-AMC, (**B**) Ac-DEVD-AMC, (**C**) Ac-YVAD-AMC, (**D**) Ac-LEHD-AMC, and (**E**) Ac-WEHD-AMC cleavage by 0.25 activity units/μL of recombinant caspase-11. Data was fitted to the Michaelis-Menten equation in GraphPad Prism and are means ± SEM of three independent experiments.

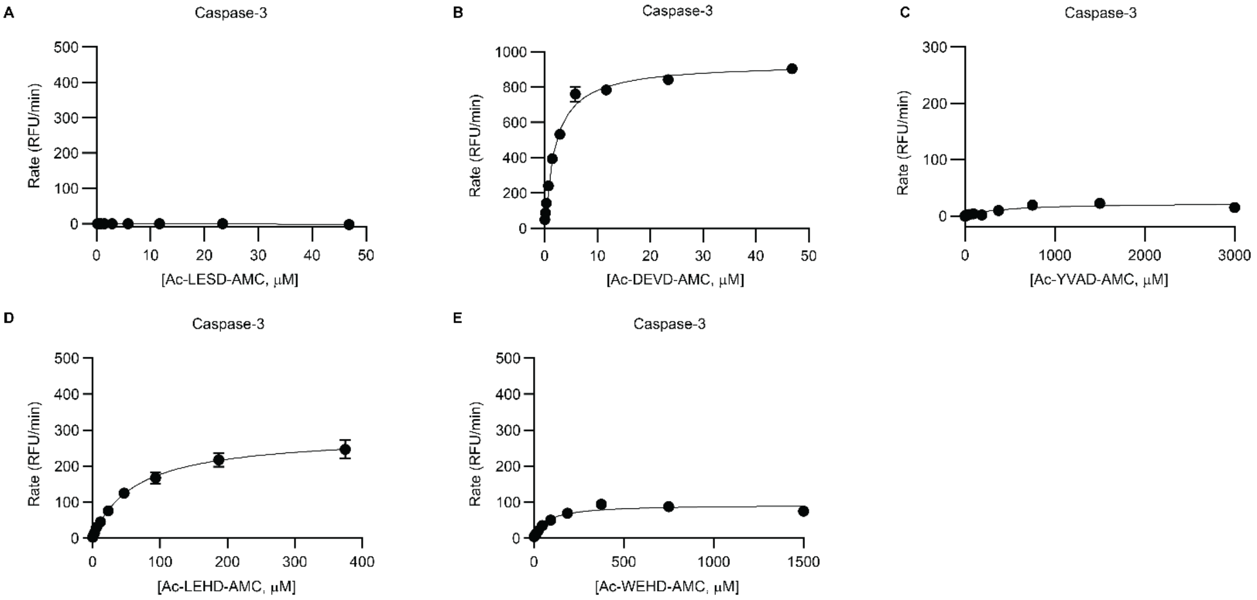


**Figure S8 (Related to Fig. 2). Kinetic characterization of the apoptotic executioner caspase-3 with tetrapeptide probes.** (**A-E**) Michaelis-Menten kinetic profiles of (**A**) Ac-LESD-AMC, (**B**) Ac-DEVD-AMC, (**C**) Ac-YVAD-AMC, (**D**) Ac-LEHD-AMC, and (**E**) Ac-WEHD-AMC cleavage by 0.25 activity units/μL of recombinant caspase-3. Data was fitted to the Michaelis-Menten equation in GraphPad Prism and are means ± SEM of three independent experiments.

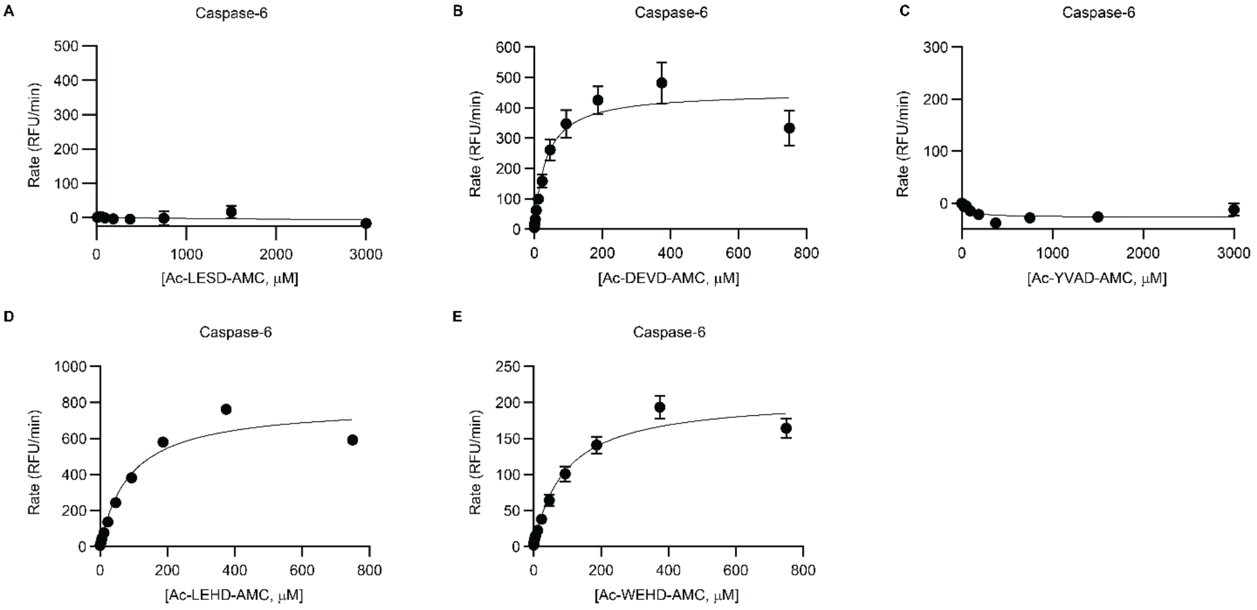


**Figure S9 (Related to Fig. 2). Kinetic characterization of the apoptotic executioner caspase-6 with tetrapeptide probes.** (**A-E**) Michaelis-Menten kinetic profiles of (**A**) Ac-LESD-AMC, (**B**) Ac-DEVD-AMC, (**C**) Ac-YVAD-AMC, (**D**) Ac-LEHD-AMC, and (**E**) Ac-WEHD-AMC cleavage by 0.25 activity units/μL of recombinant caspase-6. Data was fitted to the Michaelis-Menten equation in GraphPad Prism and are means ± SEM of three independent experiments.

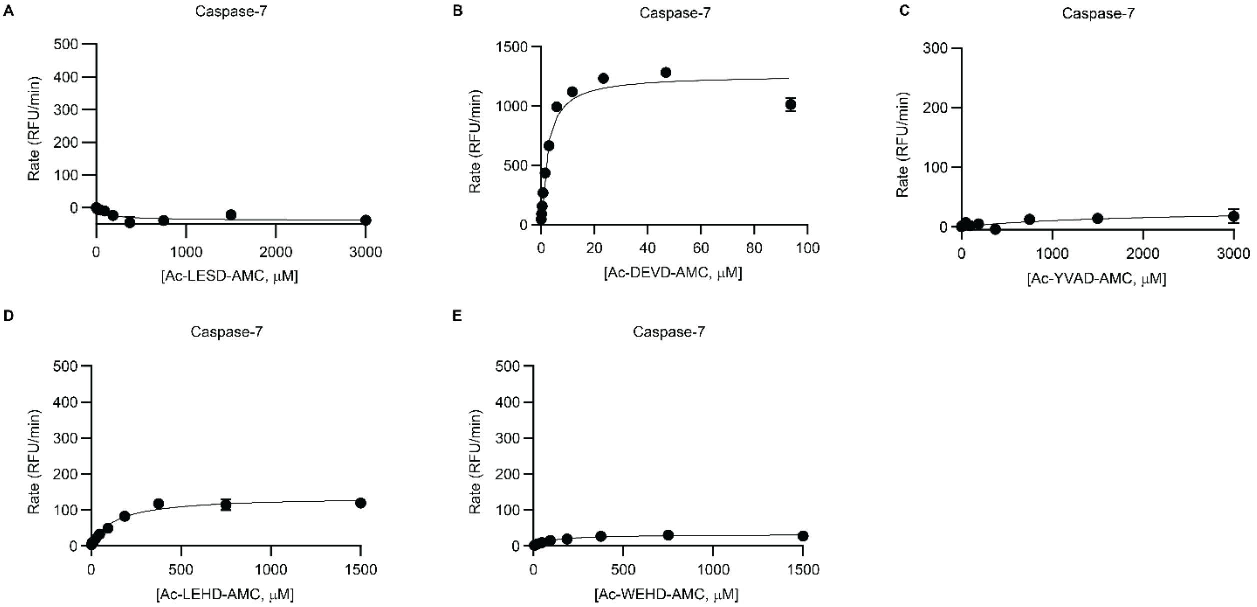


**Figure S10 (Related to Fig. 2). Kinetic characterization of the apoptotic executioner caspase-7 with tetrapeptide probes.** (**A-E**) Michaelis-Menten kinetic profiles of (**A**) Ac-LESD-AMC, (**B**) Ac-DEVD-AMC, (**C**) Ac-YVAD-AMC, (**D**) Ac-LEHD-AMC, and (**E**) Ac-WEHD-AMC cleavage by 0.25 activity units/µL of recombinant caspase-7. Data was fitted to the Michaelis-Menten equation in GraphPad Prism and are means ± SEM of three independent experiments.

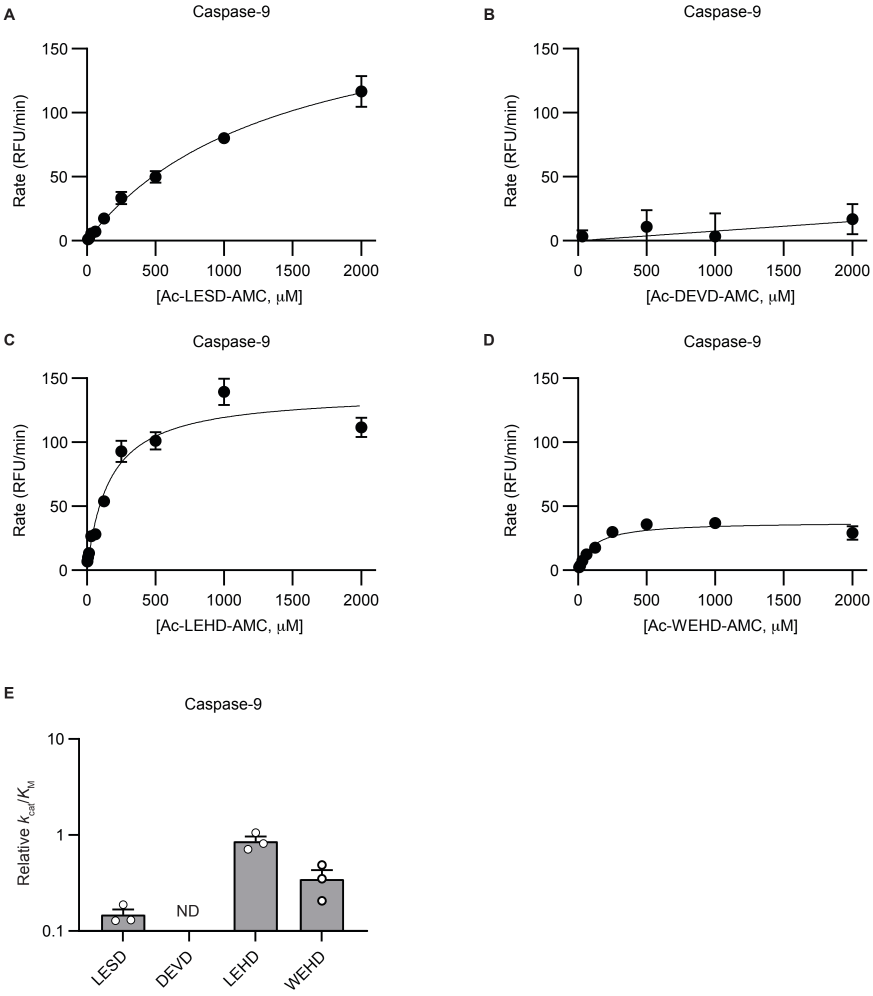


**Figure S11. Kinetic characterization of the intrinsic apoptotic initiator caspase-9 with tetrapeptide probes.** (**A-D**) Michaelis-Menten kinetic profiles of (**A**) Ac-LESD-AMC, (**B**) Ac-DEVD-AMC, (**C**) Ac-LEHD-AMC, and (**D**) Ac-WEHD-AMC cleavage by 1 activity unit/μL of recombinant caspase-9. Data was fitted to the Michaelis-Menten equation in GraphPad Prism and are means ± SEM of three independent experiments. (**E**) Relative catalytic efficiencies, represented as $k_{cat}/K_M$ of caspase-9 for each probe calculated from *in vitro* Michaelis-Menten curves.

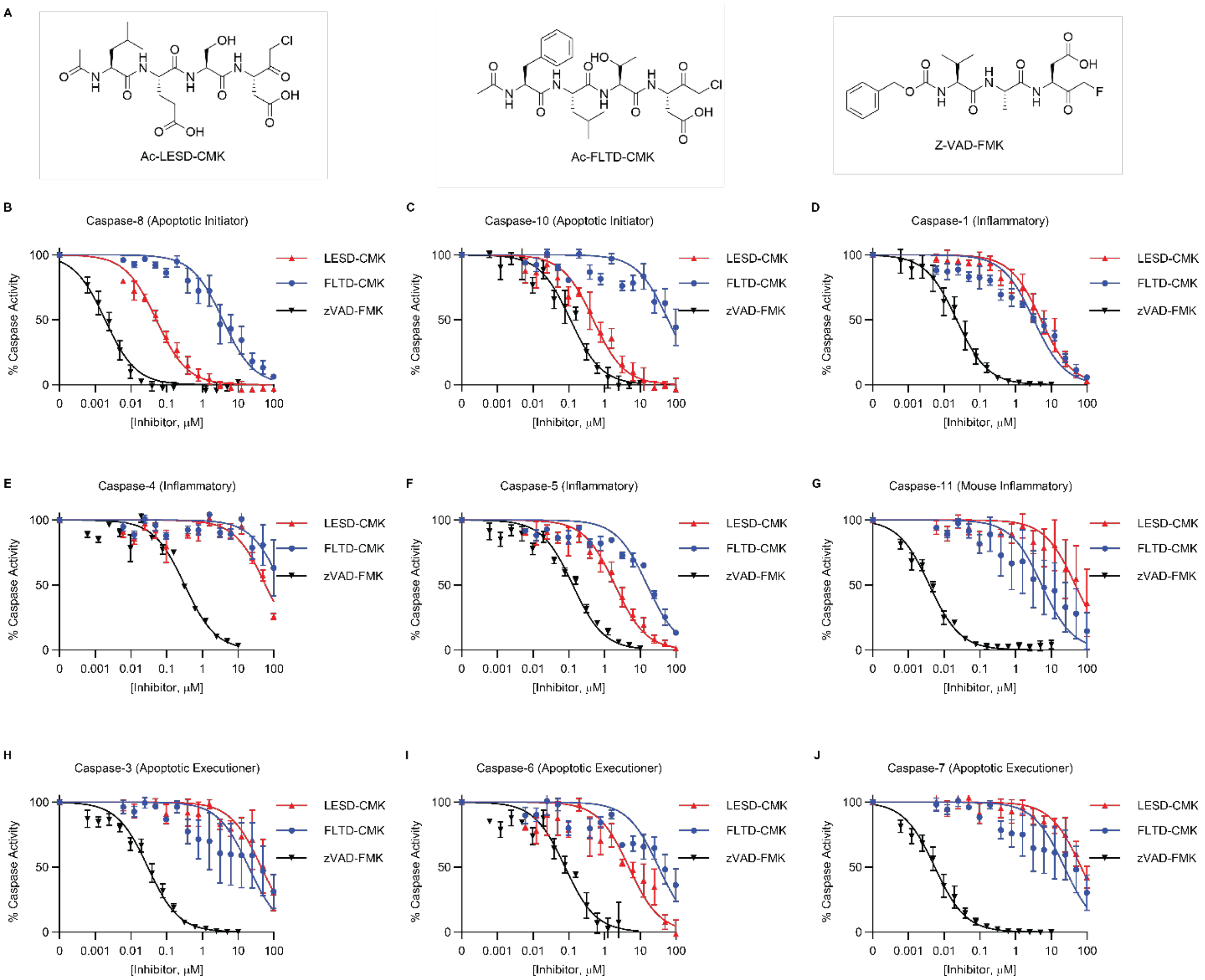


**Figure S12 (Related to Fig. 4) Selectivity profile of peptide-based caspase inhibitors continued.** Each caspase was incubated with a saturating concentration of its preferred peptide substrate, in the presence of either the Ac-FLTD-CMK, Ac-LESD-CMK, or zVAD-FMK inhibitors at indicated concentrations. Substrate cleavage rates were determined at each inhibitor concentration and normalized to the no inhibitor condition for each run. (**A**) Chemical structures of Ac-FLTD-CMK, Ac-LESD-CMK, and zVAD-FMK inhibitors. (**B**) Caspase-8 inhibition was assessed using 200 μM Ac-LEHD-AMC substrate. (**C**) Caspase-10 inhibition was assessed using 200 μM Ac-LEHD-AMC substrate. (**D**) Caspase-1 inhibition was assessed using 200 μM Ac-WEHD-AMC substrate for activity. (**E**) Caspase-4 inhibition was assessed using 1000 μM Ac-WEHD-AMC substrate. (**F**) Caspase-5 inhibition was assessed using 200 μM Ac-WEHD-AMC substrate. (**G**) Caspase-11 inhibition was assessed using 1000 μM Ac-WEHD-AMC substrate. (**H**) Caspase-3 inhibition was assessed using 100 μM Ac-DEVD-AMC substrate (25 μM of Ac-DEVD-

AMC was used for zVAD-FMK inhibition). (**I**) Caspase-6 inhibition was assessed using 100 μM Ac-DEVD-AMC substrate. (**J**) Caspase-7 inhibition was assessed using 100 μM Ac-DEVD-AMC substrate (25 μM Ac-DEVD-AMC for zVAD-FMK inhibition). Data was fitted using the [Inhibitor] vs. normalized response function in GraphPad Prism. Data are means ± SEM of three independent experiments.

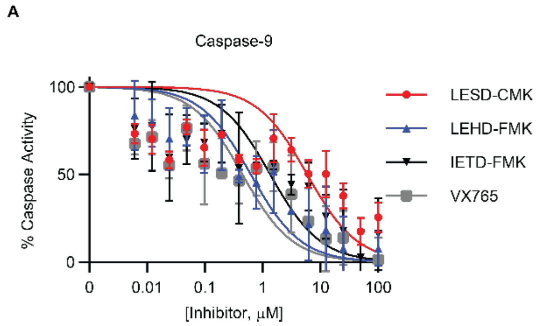


**Figure S13. Selectivity profile of peptide-based caspase inhibitors for caspase-9.** Caspase-9 was incubated at 1 activity unit/μL with a saturating concentration of Ac-LEHD-AMC (200 μM), in the presence of either the Ac-LESD-CMK, z-LEHD-FMK, z-IETD-FMK, or VX-765 inhibitors at indicated concentrations. Substrate cleavage rates were determined at each inhibitor concentration and normalized to the no inhibitor condition for each run between 20 minutes and 60 minutes. Data was fitted using the [Inhibitor] vs. normalized response function in GraphPad Prism. Data are means ± SEM of three independent experiments.
